# Receptor-based protein binding in the supramolecular network of velvet worm slime

**DOI:** 10.1101/2024.10.24.619955

**Authors:** Zhaolong Hu, Alexander Baer, Lars Hering, Ivo de Sena Oliveira, Darren C. Browne, Xue Guo, Quentin Moana Perrin, Radoslaw M. Sobota, Shawn Hoon, Georg Mayer, Matthew J. Harrington, Ali Miserez

## Abstract

The slime of velvet worms (Onychophora) is a protein-based bioadhesive that undergoes rapid, yet reversible transition from a fluid into stiff fibers used for prey capture and defense, but the mechanism by which this phase transition functions is largely unknown. Here, integrating transcriptomic and proteomic approaches with AI-guided structure predictions, we discover a group of evolutionarily conserved leucine-rich repeat (LRR) proteins in velvet worm slime that readily adopt a receptor-like, protein-binding “horseshoe” structure. Our structural predictions suggest dimerization of LRR proteins and support their interactions with conserved β-sheets-rich domains of high-molecular-weight proteins, the primary building blocks of velvet worm slime fibers. This previously unknown functional context of LRR proteins is presumably involved in reversible, receptor-based supramolecular network formation in these adhesive biofibers and provides possible new avenues for fabricating fully recyclable (bio)polymeric materials.

**Significance Statement:** Analyzing structure-function-relationships underlying reversible fiber formation in velvet worm slime may inspire avenues for the sustainable fabrication of protein-based polymeric materials. Here, we present evidence for an evolutionarily conserved mechanism of reversible fiber formation in velvet worm slime based on the receptor-like binding of fiber forming proteins by a leucine-rich repeat (LRR) protein. The structures of both protein components are highly conserved evolutionarily in the two distantly related velvet worm subgroups, indicating pervasive presence of this mechanism across species that has been maintained through the last ∼380 MY. Our results suggest that the ubiquitously occurring LRR motif—better known for its innate immunity and developmental roles—has a novel identified function in processing a biological material, which might contribute to the development of sustainable bio-inspired materials.

## Introduction

In recent decades, there has been great interest in studying living organisms producing protein-based structural materials that function outside their bodies—such as spiders and mussels—as bioinspiration to fabricate polymeric materials in a more sustainable way (1, 2). In this regard, the slime of the velvet worms (Onychophora) has emerged as an intriguing model system (3) for its notable ability to transform instantly from an adhesive fluid into rigid fibers initiated by a simple mechanical trigger (3, 4). Unlike fibers produced by spiders and mussels, however, the mechanically activated formation of fibers from velvet worm slime occurs entirely outside the animal’s body and thus without specific regulatory input by the organism. Those fibers, with a stiffness in the dry state comparable to nylon, are capable of dissolving back into their biomolecular precursors under aqueous conditions. Remarkably, new fibers can be drawn from this solution, indicating that instructions for mechanoresponsive fiber self-assembly are encoded within the biomacromolecular building blocks themselves (3). Both the mechanically triggered self-assembly of a protein solution into fibers and the recyclability of these fibers have motivated detailed investigations of this natural adhesive (1, 3).

Velvet worms comprise a phylogenetically ancient group of terrestrial invertebrates, which are organized in two geographically distinct major subgroups, Peripatidae and Peripatopsidae that diverged over 380 MY (5-8). As a conserved feature of all velvet worms, they capture prey and defend themselves by rapidly ejecting two jets of fluid slime from specialized papillae on each side of their head. Once the victim attempts to escape, the slime first transforms into a sticky viscoelastic gel and eventually into stiff and glassy fibers as it dries (7-13). Across different species studied, velvet worm slime consists of ∼90% water. The dry weight contains at least 50% proteins organized roughly into three classes of molecular weight (MW) ranging from 10–280 kDa. More recently, encapsulated phosphate and carbonate salts are found to constitute a significant 12–6 wt% of the initial mass (14). Carbohydrates occur only in minute quantities (∼2%), mostly linked to proteins, and lipids are even less abundant. The remaining rest of dry matter is unknown up to this point (4, 7, 13, 15-18). Recent nano-structural investigations suggest that high-MW proline (Pro)-rich proteins (>200 kDa), which are presumably the key structural element of the fibers, exist freely in solution as monomers and larger MW complexes (15, 16, 19).

Currently, the full-length sequences of high-MW slime proteins have been identified only in a single undescribed species from Singapore (presumably peripatid *Eoperipatus* sp.), with conformational predictions of a mixture of random coil structure and several small β-sheet rich domains (19). Prior research has also shown the presence of mid-MW proteins (75–110 kDa) that have not yet been characterized regarding their sequence, structure and possible functional role in the slime. It has only been hypothesized that, together with lipids, they form the characteristic monodisperse nanoglobules that were previously observed in slime from several species (3, 16, 19, 20). Here, we focus specifically on these understudied, yet prominent mid-MW slime proteins.

Although there have been some insights into the physical and chemical principles underlying reversible fiber formation of velvet worm slime including the role of electrostatic interactions, disulfide bonding, and protein conformational transitions (4, 19-21), the very fragmentary sequence data impede further progress. Indeed, this information is essential to understand the key structure-function relationships and interactions of specific proteins which underly the ∼380-MY conserved mechanisms of fiber formation in velvet worms (22). Particularly, the co-conservation of high- and mid-MW proteins across evolutionarily distant velvet worm species suggests a highly conserved mechanism of protein assembly involving interactions between specific proteins (15, 16). Here, we take a combined RNA-sequencing and proteomics approach coupled to AI-guided structural predictions of high- and mid-MW proteins from three distantly related species representing both major onychophoran subgroups. Our results reveal receptor-like mid-MW slime proteins that are conserved across all three species and feature the characteristic “horseshoe” structure of ligand-binding domains in leucine-rich repeat (LRR) containing proteins. Our predictions indicate the likelihood of homo- and hetero-dimerization of the LRR motifs in the mid-MW proteins, and that conserved β-sheet-rich domains of the high-MW proteins are possible binding partners of these LRR proteins. These findings suggest that reversible receptor-based interactions between LRR and high-MW proteins are involved in the rapid fiber formation process in velvet worm slime that has been conserved for nearly 400 million years, providing potential molecular guidelines for sustainable fabrication of recyclable polymeric materials.

## Results

### Molecular weight distribution of slime proteins

The MW distribution of slime proteins in three species representing both major velvet worm subgroups, namely *Euperipatoides rowelli* (Peripatopsidae, Australia), *Eoperipatus* sp. (Peripatidae, Singapore) and *Epiperipatus* cf. *barbadensis* (Peripatidae, Barbados) (Fig. 1; distribution and analysis of phylogenetic relationships are shown in *SI Appendix*, Fig. S1 and Supporting Information Text), was first analyzed using SDS-polyacrylamide gel electrophoresis (SDS-PAGE). In line with previous reports (15-19, 23), protein bands in all species are located in three main regions of MW (high > 230 kDa; mid: 70–110 kDa; and low-MW < 70 kDa). The high-MW proteins are most abundant and consist of multimer complexes linked by disulfide bonds based on changes in band mobility upon disulfide bond reduction. Although differences in the band profiles are apparent (mostly between representatives of Peripatidae and Peripatopsidae), the existence of both the high- and mid-MW proteins is conserved between all species. In contrast, low-MW proteins show a large interspecific variability even within the same velvet worm subgroup (Fig. 1 and *SI Appendix*, Fig. S2). Given the distant phylogenetic relationship of the three species studied, the simultaneous conservation of high- and mid-MW proteins over at least 380 MY likely indicates their important function in velvet worm slime, motivating further investigations of protein sequences and conformational features, as well as possible interactions between these proteins.

**Figure 1.**
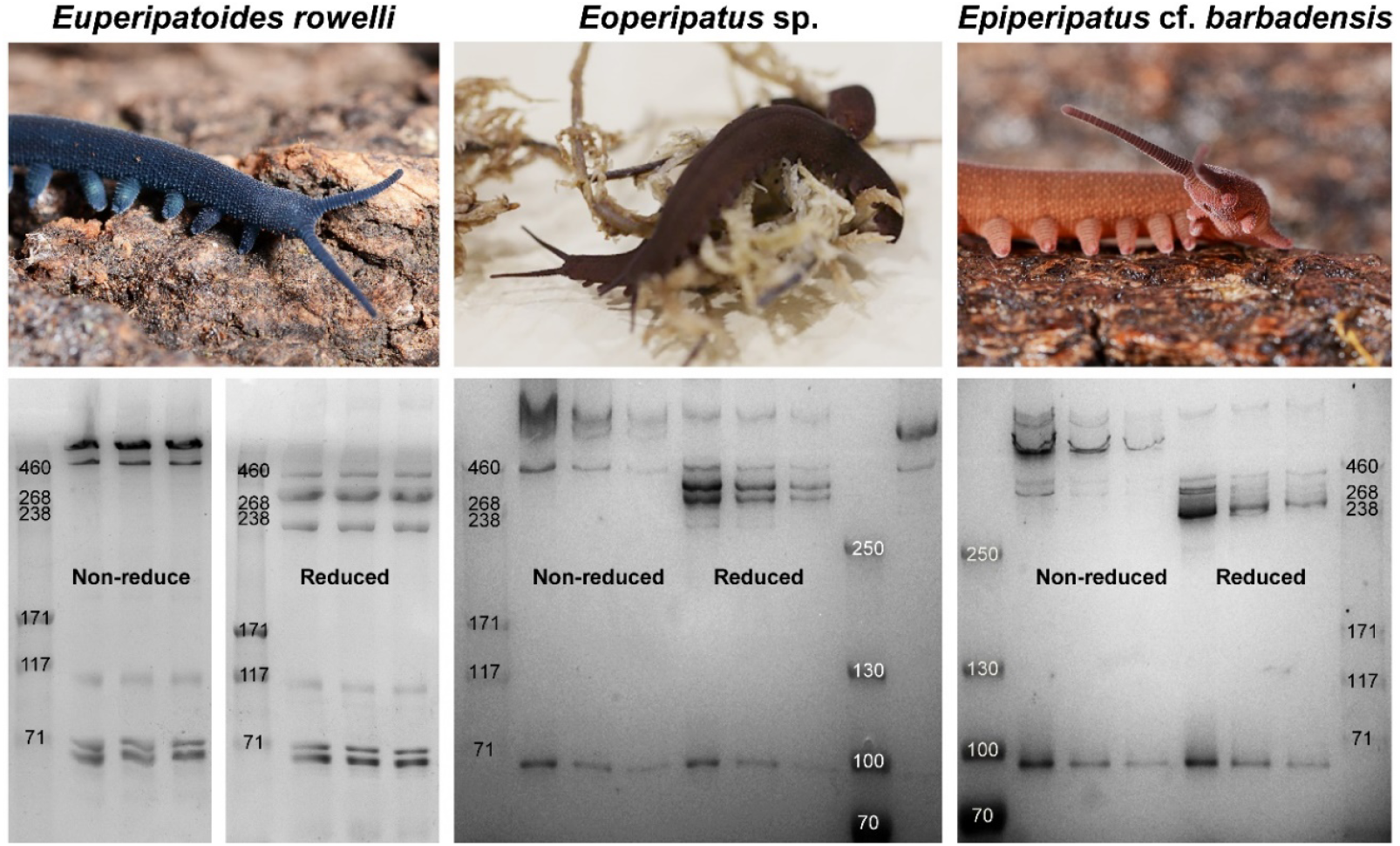
Representatives of velvet worm species studied and their typical molecular weight distribution of slime proteins higher than 70 kDa. (Above) *Euperipatoides rowelli* from Australia (Peripatopsidae), *Eoperipatus* sp. from Singapore and *Epiperipatus* cf. *barbadensis* from Barbados (Peripatidae) from left to right. (Below) Denaturing polyacrylamide electrophoresis (SDS-PAGE) of slime proteins using a gradient 5% polyacrylamide gel under non-reducing and reducing conditions. Low-MW region is shown in *SI Appendix*, Fig. S2. Gel images of *Eu. rowelli* are adopted from Baer *et al*. (16) under the CC-BY-NC license.

### Identification of leucine rich (LRR) repeat proteins and their structural similarities to LRR-receptor proteins

Proteins in the mid-MW range were previously reported in velvet worm slime from different species (15, 16), but their molecular, structural, and functional characteristics have remained unexamined due to the lack of available sequence data. Using transcriptomic and proteomic analysis of the protein bands between 70 kDa to ∼110 kDa (Fig. 1 and *SI Appendix*, Fig. S2), we identified 3–4 protein sequences in each species. Each transcript is predicted to have both a eukaryotic signal peptide and a stop codon, indicating sequence completeness (*SI Appendix*, Table S1). The identified proteins can be clearly distinguished into two categories based on sequence and structure similarities. The LRR proteins comprise most mid-MW proteins and are the focus of this study. For the less abundant proline-rich mid-MW proteins, two transcripts were identified in *Eoperipatus* sp. They are similar to those previously reported for *Eu. rowelli* [HM217028, HM217029 (18)] and resemble a shorter version of the high-MW proteins, based on the high content of Pro and charged amino acids (*SI Appendix*, Tables S1–2), as well as predicted structural similarities (*SI Appendix*, Figs. S4D and S9).

The mid-MW proteins of the predominant LRR group feature high molar content of leucine (Leu, L)/isoleucine (Ile, I), and both positively and negatively charged residues (each over 20% of total amino acids) (*SI Appendix*, Figs. S3A–B, mid-MW protein sequences and Table S2). The number of identified transcripts differs between the species, which also include potential isoforms of the same proteins referred to as variants, *i*.*e*. ER_PM1-var1-3, EB_PM1-var1-3, and ES_PM1 (*SI Appendix*, Fig. S3C). Most notably, positioning of Leu and Ile in these proteins and across all species is highly conserved in the LRR motifs, with a consistent length of 24 amino acids (Fig. 2A and *SI Appendix*, Table S3). Each of these repeats resemble the hallmark 11-residue sequence motif LxxLxLxxNxL (N is asparagine (Asn), x is any amino acid) of the well-described “typical” LRR subfamily (InterPro entry IPR003591) (24-27). For LRR proteins in velvet worm slime, we identified the consensus motif sequence: LxxLxxLxLxxNxIxxIxxxxFxx (where F is phenylalanine, Phe). When compared across the species, the Ile at positions +14 and +17 are highly conserved in *Eu. rowelli* but there is an equal observed occurrences of Leu at the same positions in the other two species. The observed occurrences of Phe at position +22 is lower in *Epiperipatus* cf. *barbadensis* (Fig. 2A). BLAST searches against protein databases suggest that velvet worm LRR proteins show the strongest homology with LRR-containing proteins featuring receptor-like binding domains, including insulin-like growth factor-binding and chaoptin-like proteins, and LRR-containing protein 15 (*SI Appendix*, Table S4).

**Figure 2.**
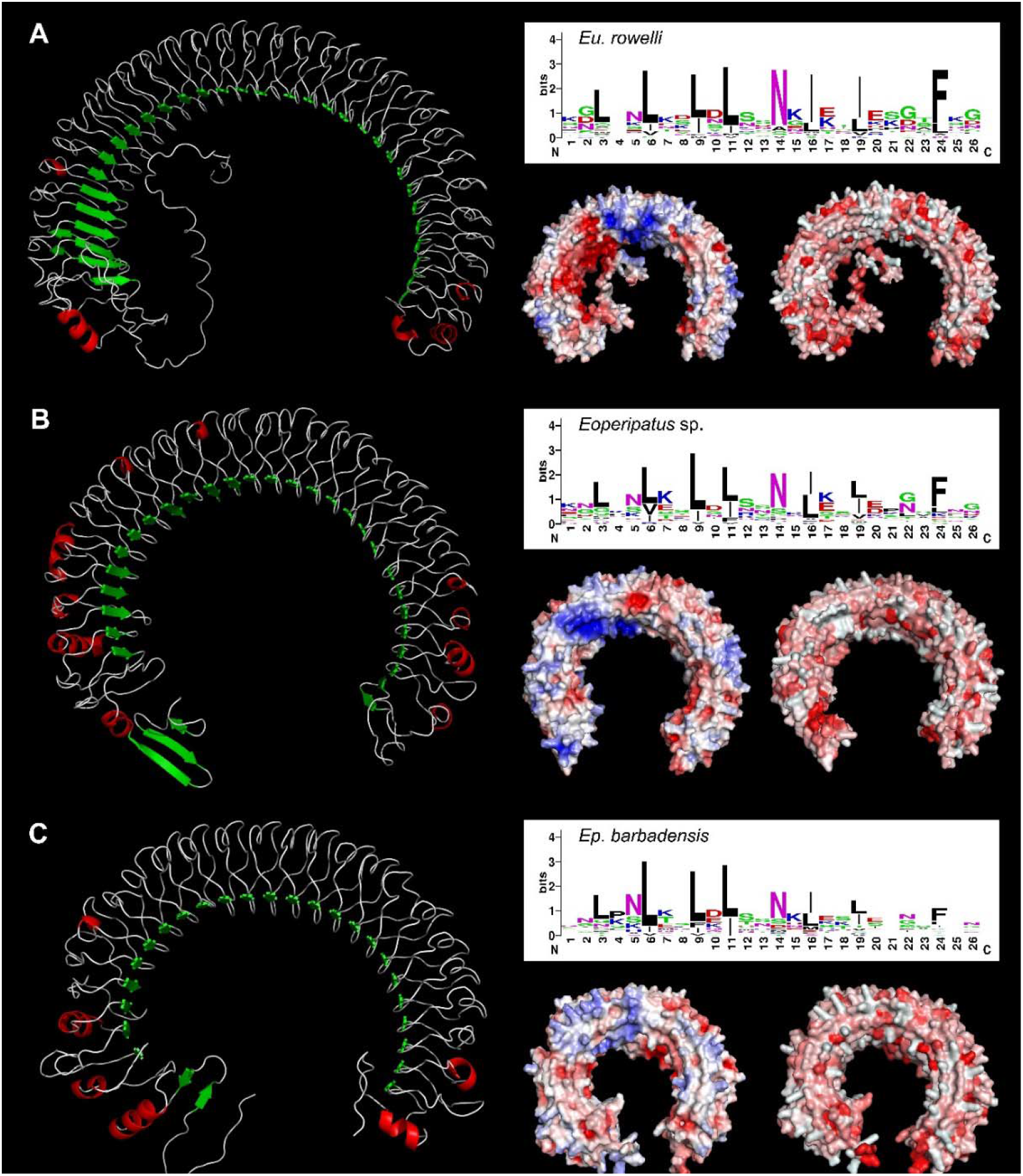
Structural analysis of representative mid-MW LRR proteins in three species studied. (A) *Eu. rowelli* (B) *Eoperipatus* sp. (C) *Ep*. cf. *barbadensis*. (Left) AlphaFold3 predicted structures with annotated β-sheets highlighted in green and α-helices in red. (Right) Sequence logo plot of interspecific conserved Leu, Ile and Asp identified in each of the 24 LRR loops. Surface charges at native pH 5.2 (positively charged in blue, negatively charged in red) and surface hydrophobicity (hydrophilic in white, hydrophobic in red) are displayed in the bottom left and right panels, respectively.

LRR motifs have been identified in highly diverse proteins in most animals from cnidarians to vertebrates, as well as in plants and bacteria (28, 29). Although existing LRR domains might be phylogenetically and functionally unrelated, they provide a conserved structure featuring an overall non-globular arc-like, “horseshoe” or “solenoid” shape constructed from solvent-exposed parallel β-sheets, which provides accessible hydrophobic binding sites directly involved in binding ligands such as proteins, nucleic acids, lipids, and lipoproteins (24-26, 29-31). Functions of LRR domains range from cell adhesion and signaling, extracellular matrix assembly, neuronal development and neural functions, RNA processing, adhesion and invasion of pathogenic bacteria to host cells, disease resistance and pathogen recognition in plants and immune response (24, 26-29).

Similar to extracellular, ligand binding domains of LRR receptor proteins such as Toll receptors in *Drosophila* and *Aedes* (24, 30, 31), AlphaFold3-based structural predictions show that LRR proteins of all velvet worm species fold into the characteristic “horseshoe” structures (Figs. 2A–C, *SI Appendix*, Movie M1). The overall sequence homology among the LRR proteins identified in slime of three species is rather low (*SI Appendix*, Fig. S3C). However, the protein structure is evolutionarily conserved due to highly conserved Leu motifs which indicates that these proteins might play a key role in the slime. The protein structure is stabilized by a regular arrangement of 21–34 loops that feature alternating hydrophobic and hydrophilic side chains. The hydrophobic residues are mostly oriented into the inner core of the “horseshoe” shape while the hydrophilic residues are located on the outer surface, likely responsible for the water-soluble character of these proteins (Fig. 2A-C, right panels and *SI Appendix*, Fig. S5). On the concave side, the loops show flattened β-strands of three hydrophobic amino acids (five longer β-strands of 6–8 amino acids occur at the N-terminus), which are connected by five hydrogen bonds into a curved, parallel β-sheet consisting of over 30 individual β-strands (*SI Appendix*, Fig. S6). These predictions are consistent with previous wide-angle X-ray diffraction (WAXD) studies that indicated the presence of narrow, elongated β-sheets (20). Specifically, Scherrer analysis of the diffraction peak widths predicted β-sheet crystallites of 10.5 nm by 1.3 nm, approximately corresponding to a sheet comprising 22 strands, each 3–4 amino acids long, which is remarkably consistent with the AlphaFold3 predicted structure, within the error of the measurements.

*In vitro* evidence for β-sheet conformation was obtained by Circular Dichroism (CD) spectroscopy on synthesized peptides derived from single Leu-rich loops of LRR mid-MW proteins in velvet worm slime. The data show that the peptides self-assemble into β-sheet structures—independent of the presence of lipids—that are stable over time (*SI Appendix*, Fig. S7). The outer surface on the convex side adopts non-canonical random-coil structures, short β-strands and very few short α-helix-turns (Fig. 2). The loops are fan-like without interconnecting bonds, which may provide binding sites for interlocking interactions not only with other slime proteins but also other macromolecules. Only a few continuous hydrogen bonds run perpendicular to the polypeptide backbone in order to provide stability within the loop structure (Fig. 2, *SI Appendix* Fig. S6) (24). The N- and C-termini in all LRR proteins show concentrated solvent exposed hydrophobic residues (*SI Appendix*, Fig. S5). Further, the N-terminus of LRR proteins are capped by a non-repetitive around 150 amino acids long domain, which contains three interspecifically conserved Cys residues (*SI Appendix*, Fig. S3C) that might provide stability and solubility to the hydrophobic LRR (32, 33).

### Identification of high-MW proteins that provide possible binding partners to LRR proteins

Compelling similarities of velvet worm LRR proteins to ligand-binding LRR proteins such as Toll receptors, as well as conserved LRR motifs and structures, motivated us to search for possible binding partners. In *Drosophila* and *Aedes* innate immunity and cell signaling, the Toll receptor is activated by binding to the endogenous ligand Spaetzle (Spz) (29). Spz is a cytokine intermediate consisting of two protein chains forming two opposing β-sheets, each comprising four antiparallel β-strands (30). Interestingly, a very similar tertiary structure was previously identified in specific domains of high-MW slime proteins of *Eoperipatus* sp. from Singapore. Drawing analogy to Toll-Spz binding, these domains could present a binding partner for the mid-MW LRR protein (19). To assess whether these distinctive protein folds in the high-MW are conserved in the other species, we then undertook transcriptomic and proteomic analyses of the SDS gel protein bands larger than 200 kDa for all three species.

From the high-MW region of SDS-gels, sequences of two predominant proteins were obtained for each species (Fig. 3A and *SI Appendix*, Tables S5–6) (at least one further high-MW transcript occurs in the slime according to our transcriptome and SDS gel data; however, this was not analyzed further here). Completeness of encoding genes was verified, leading to corresponding single nucleotide and insertion/deletion (indel) polymorphisms (*SI Appendix*, Table S7). Only the N-terminal region for ER_PH2 was not identified. Comparisons of high-MW proteins in *Eu. rowelli* (ER_PH1 and ER_PH2) and *Epiperipatus* cf. *barbadensis* (EB_PH1 and EB_PH2) with previously verified sequences ES_PH1 and ES_PH2 from *Eoperipatus* sp. (19) reveal major overall similarities across the species in terms of: (i) the general amino acid composition; (ii) distinct N- and C-terminal domains; (iii) largely disordered and repetitive core; and (iv) high degree of post-translational modifications (PTMs) including hydroxylation of Pro, glycosylation and phosphorylation, suggesting evolutionarily conserved features in high-MW proteins in velvet worm slime over a span of at least 380 million years (Fig. 3A and *SI Appendix*, Figs. S8–9).

**Figure 3.**
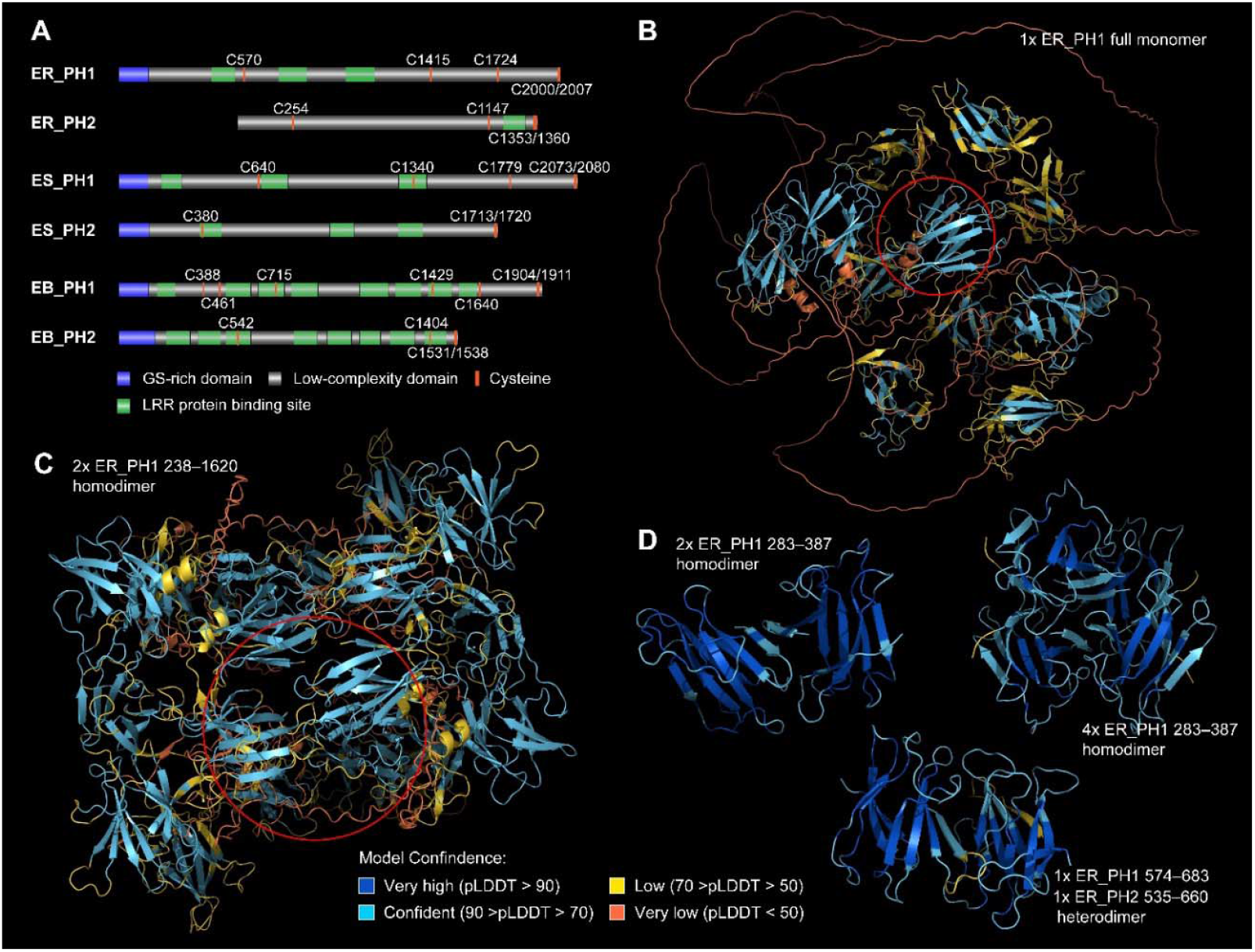
Domain architecture, predicted structure, and interactions of high-MW proteins via β-sheet-rich domains. (A) Identified high-MW proteins in slimes of three species (ER = *Eu. rowelli*; ES = *Eoperipatus* sp.; EB = *Ep*. cf. *barbadensis*). The lengths of sequences are proportional to reconstructed sizes. Blue and grey bars represent GS-rich and the low-complex core domain, respectively. Green regions highlight predicted binding hot spots for LRR proteins. Orange lines mark the position of Cys residues. (B) Structure of the high-MW protein PH1 in *Eu. rowelli*. (C) Predicted interactions between two high-MW proteins (sequences without N- and C-termini). (D) Predicted interactions between two and multiple β-sheet-rich domains of ER_PH1 (top), and interactions of β-sheet-rich domains of high-MW proteins PH1 and PH2 (bottom). Red circles highlight β-sheet-rich domains and their intra- and intermolecular interactions. Model confidence values (pLDDT) are indicated by coloration of predicted structures.

In line with previous reports from small-angle neutron scattering (SANS) (16), structure predictions indicate a globular tertiary structure of the high-MW proteins in all species studied (Fig. 3B). While the N- and C-termini lack any secondary structure and abundant Pro lead to an overall largely disordered protein structure, the core of the protein features localized domains with well-defined conformations (Figs. 3B–C, *SI Appendix*, Fig. S9 and Movie M2). Indeed, the AI-guided structural predictions of these proteins using AlphaFold3 confirm regularly-spaced domains of antiparallel β-strands 4–8 amino acids long that are organized into twisted arrangements of two β-sheets, each consisting of four or less abundant three β-strands (Fig. 3B and *SI Appendix*, Figs. S4B–D). In contrast to previous suggestions of electrostatic stabilized β-sheets in velvet worm slime (20), our data clearly show that these antiparallel β-strands exclusively occur in hydrophobic regions (*SI Appendix*, Fig. S10). AlphaFold3 predictions of the core sequence of high-MW proteins and various separated β-sheet containing sequence regions reveal with high confidence that β-sheet domains can interact with each other (Figs. 3C–D). These results suggest the presence of isolated globular high-MW proteins in the non-agitated slime that are stabilized by intramolecular interaction of hydrophobic β-sheet domains, which prevents premature aggregation with other molecules. However, when brought in proximity by drying or shear forces, the β-sheet domains are exposed and may readily interact with those of other high-MW proteins, which provides an unspecific and fast, yet reversible aggregation mechanism.

### AI-guided predictions of interactions between LRR proteins and β-sheet-rich domains of high-MW proteins and LRR dimerization

Ligand binding by LRR receptor proteins is reported in numerous studies, whereby protein ligands with radii in the range of the arc of the horseshoe structure are largely surrounded by the concave surface of the parallel β-sheets of the LRR domain, maximizing the number of protein-protein contacts (24). In case of the Toll/Spz complex, reversible ligand binding typically occurs at the N-terminal β-sheet domain of the concave side, resulting in receptor-ligand heterodimers (30, 31). These interactions achieve equilibrium binding constants in the range of *K*_D_ = 82–92 nM, indicating a high affinity, yet still dynamic binding pair (34).

Notably, β-sheet-rich domains of high-MW slime proteins bear some general similarities to the Spz ligand in one aspect, namely they also contain six antiparallel hydrophobic β-strands arranged into a twisted β-sheet-rich domain. Thus, we decided to investigate possible interactions between LRR proteins and the β-sheet-rich domains of high-MW proteins. AlphaFold3 confidently predicts protein interactions between LRR proteins and isolated single and multiple β-sheet-rich domains of both PH1 and PH2 high-MW proteins, with binding sites within the horseshoe arc (Figs. 2A–C and Fig. 3A). Furthermore, for β-sheet domains of the non-LRR mid-MW proteins in *Eoperipatus* sp. and those previously reported for *Eu. rowelli* [P2a, P2b (18)], binding via LRR proteins is also predicted with sufficient confidence (Fig. 3D). This also applies to the N-or C-terminal regions of the high-MW proteins. Instead, our results suggest that the C-termini of the high-MW proteins interact preferably with each other, which is in line with previous reports of the formation of high-MW multi-protein complexes (16, 19). The origin for this preference might be the interspecifically highly conserved C-terminal motif that consists of regularly arranged charged and hydrophobic residues and well-aligned Cys residues positioned exactly seven amino acids apart (Cys_30_ and Cys_37_, i/i+7 pattern, *SI Appendix* Fig. S11B). We conclude that LRR proteins in the velvet worm slime preferably bind specific β-sheet-rich regions of the high- and mid-MW slime proteins.

For several LRR receptor-like proteins, mono-as well as multi-dimerization has been reported to occur through various interaction domains, including the concave surface, the ascending side, and at both terminal regions (24, 31). AlphaFold3 predictions confirm the dimerization of velvet worm LRR proteins in versatile orientation, which also includes head-to-tail interlocking and interactions on both the concave and the convex surfaces (Fig. 4B). Notably, predictions of multiple LRR protein sequences with (i) several isolated β-sheet rich domains of high-MW; (ii) partial sequences of high-MW proteins comprising all β-sheet rich domains (*e*.*g*. core part without N- and C-termini); and (iii) full sequences of non-LRR mid-MW proteins [ES_PM2, previously published P2a and P2b from *Eu. rowelli*, HM217028, HM217029 (18)] confidently indicate that dimerized LRR proteins are capable to bind high-MW and non-LRR mid-MW proteins through interaction between hydrophobic β-sheets (Figs. 4C–E).

**Figure 4.**
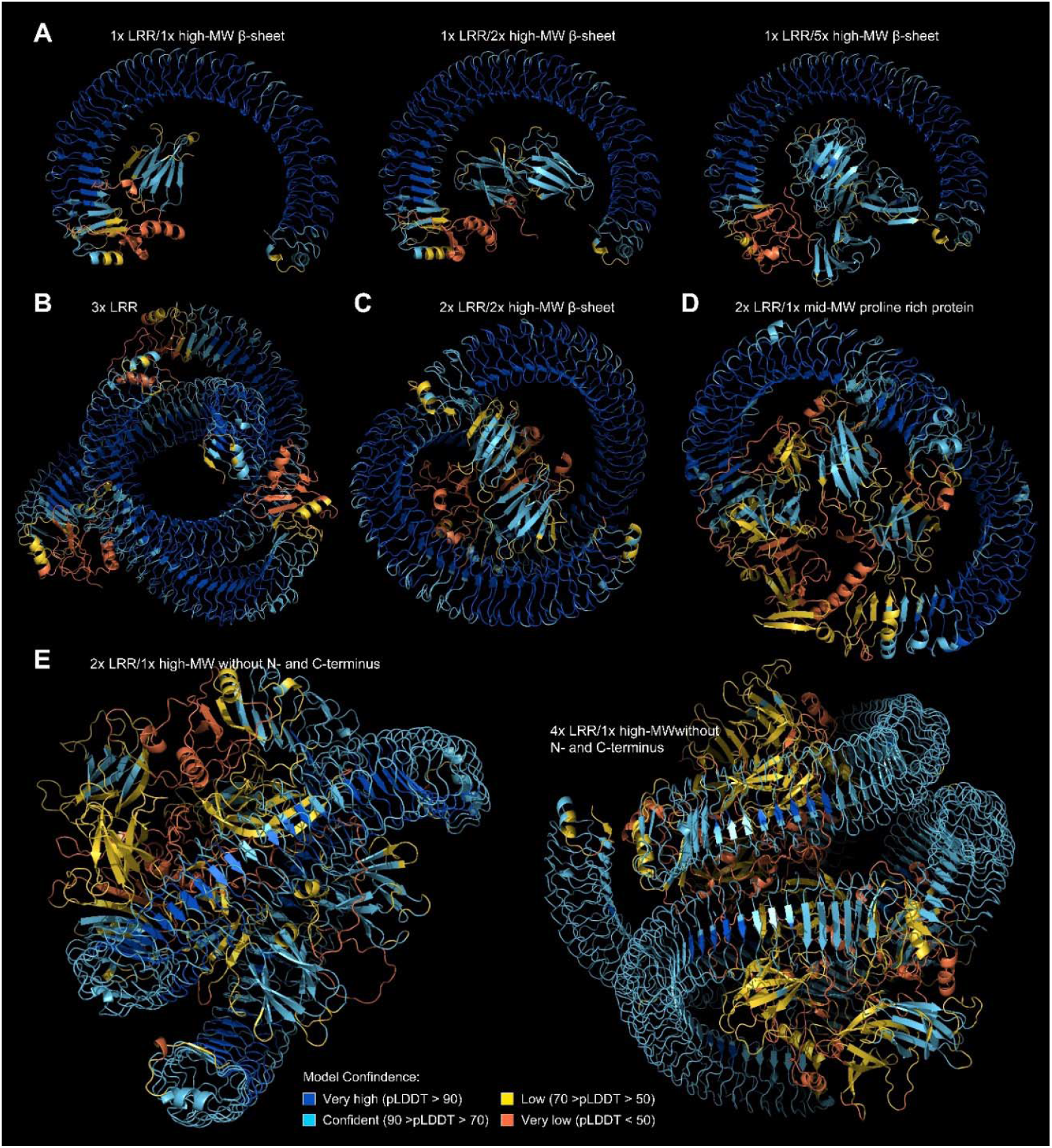
*In silico* evidence that LRR proteins bind β-sheet-rich domains of high-MW and non LRR mid-MW slime proteins. (A) LRR proteins (ER_PM1-var1) are predicted to bind one up to multiple isolated β-sheet-rich domains of both high-MW proteins ER_PH1 and ER_PH2 with high confidence. (B) Predicted homo-multimerization of three LRR proteins (ER_PM1-var1) showing head-to-tail oriented interlocking and binding via the concave and convex sides. (C) Predicted interaction between a LRR protein dimer and two β-sheet-rich domains of the high-MW slime protein. (D) Predicted interaction between LRR protein dimer (ES_PM1) and full sequence of non-LRR mid-MW protein (ES_PM2). (E) Prediction of two and four LRR-proteins (ER_PM1-var1) binding β-sheet-rich domains from partial sequence of a high-MW protein (ER_PH1 without N- and C-termini).

In further support, multiple ligand binding by dimerized receptors is reported from head-to-tail oriented homodimers of Toll-like receptor proteins in the mosquito *Aedes aegypti*, which are able to reversibly bind two Spz ligands (31). Spz typically binds the β-sheet of the concave N-terminal region; however, a second Spz ligand is predicted to bind at the convex dimerization site of the two receptor proteins, forming a very stable 2:2 heterodimer (31). These results from *in silico* analyses of the key structural building blocks in velvet worm slime support potential mechanisms by which the Pro rich high-MW proteins can form specific, yet reversible intermolecular interactions with the mid-MW LRR proteins, which could conceivably play a key role in protein aggregation and thus in the reversible formation of fibers.

## Discussion

LRR receptor-based ligand binding is a well described principle in nearly all groups of organisms across bacteria, plants and animals, functioning in cell adhesion, development and innate immunity (24, 26-31). Here, we have identified free extracellular LRR receptor-like proteins and putative ligand domains in velvet worm slime employed in what appears to be a distinctive functional context—macroscopic, extra-corporeal material self-assembly. Although further experimental support is needed, it is tempting to speculate that such dynamic ligand-receptor based interactions might regulate the reversible mechanoresponsive fiber formation in velvet worm slime—a hypothesis that is supported by the fact that key structural elements in these two protein families have been preserved for nearly 400 million years. At the most basic level, one could envision that velvet worm slime fiber formation occurs when LRR receptor domains of the mid-MW proteins bind to the conserved β-sheet-rich domains of high-MW proteins, creating a large supramolecular network structure. This would naturally require the creation of network nodes, in which either one LRR receptor binds multiple ligands from at least two different high-MW protein chains, or dimerized LRR domains each bind ligands on different high-MW protein chains—scenarios that are both supported by our ligand docking models.

While our analysis clearly suggests that interactions between LRR proteins and β-sheet-rich domains of high-MW proteins are likely involved in fiber formation, the question of the mechanoresponsive and reversible nature of fiber formation is more complex. Indeed, fiber formation is a multi-step process in which fluid slime must be mechanically activated to form a viscoelastic gel phase through shearing before it can be drawn into stiff and glassy fibers via drawing. Assuming that network formation depends on receptor-ligand interactions between high- and mid-MW proteins, it stands to reason that the relevant domains are segregated from one another in the fluid state. Our models predicted that the β-sheet-rich domains in the high-MW proteins tend to cluster in the protein core while it is proposed that the mid-MW proteins are partitioned within the slime nanoglobules (16), thus preventing the proteins from interacting. However, shear forces could conceivably function to release mid-MW proteins from the nanoglobules as well as elongate the high-MW proteins, therefore exposing cryptic ligands and favoring the formation of a network based on receptor-ligand binding interactions, eventually leading to a hydrogel phase that can subsequently be drawn into a fiber and dried. Notably, if the gel phase is not dried, it will eventually relax back into the fluid slime phase (3).

Based on our current findings, we posit that mechanical shear takes the system out of the equilibrium fluid phase, creating a temporarily stable gel phase via formation of a network of dynamic receptor-ligand interactions. However, if the concentrations of receptors and ligands are kept constant (*i*.*e*., the gel is not dried), it will relax back to the lowest energy state determined by the equilibrium binding constants, which is the fluid slime phase. This would also explain why fibers can be solubilized in excess water, which is reported for *Eu. rowelli* (3) and observed in *Eoperipatus* sp. (19). On the other hand, elongational drawing of fibers could favor stronger network formation by exposing more ligands and receptors via mechanical shear. This mechanical trigger could also increase the surface area and induce drying, which would further enhance the concentration of receptors and ligands and shift the equilibrium toward stronger interactions. Nevertheless, although experimental evidence for multi-dimerization and their specific function in the fiber formation still needs to be obtained, receptor-based protein assembly represents an intriguing new avenue towards the fabrication of sustainable biomimetic polymers and materials.

## Materials and Methods

### Sample collection and preparation

#### Collection of specimens

Specimens of *Euperipatoides rowelli* Reid, 1996 were collected in Australia (New South Wales, Tallaganda State Forest, 35°26’S, 149°33’E). Specimens of *Epiperipatus* cf. *barbadensis* (Froehlich, 1962) were collected in Barbados (Hope Road Gully, St. George, 13°08’51’’N, 59°33’12’’W). Specimens of undescribed *Eoperipatus* sp. were collected in a local secondary forest in Singapore near the coast of the island Pulau Ubin (1°24’04.3”N 103°58’04.7”E).

#### Export and collection permits

Specimens of *Eu. rowelli* were collected and exported under the permit numbers: SL101720/2016, issued by NSW National Parks & Wildlife Service (Australia), and PWS2016-AU-001023, issued by Department of Sustainability, Environment, Water, Population, and Communities (Australia). Specimens of *Eoperipatus* sp. were collected under the Permit No NP/RP19-037 from the National Parks Board, Singapore. Specimens of *Epiperipatus* cf. *barbadensis* were collected and exported under the permit 8434/56/1, issued by the Ministry of Environment and National Beautification, Green and Blue Economy, Barbados.

#### Animal maintenance, and collection and preparation of slime samples

Specimens of *Eoperipatus* sp. and *Epiperipatus* cf. *barbadensis* were kept at a temperature of 20–26 °C, those of *Eu. rowelli* at 18 °C. The worms were kept on moist Sphagnum moss or peat and were fed with crickets. The slime collection was performed, by stimulating the specimens to shoot into a vail or the ejection of slime was directed to a pipette tip and allowed to dry. Solid dry flakes were redissolved in deionized water over 10h–12hours with gentle agitation to prepare aqueous slime solution and stored at 4 °C until use.

#### Sample preparation and SDS-PAGE

Protein concentrations in collected slime sample were detected using a Qubit fluorometer (Thermo Fisher, United State) with protein broad range assay kits. About 100 μg of protein with 2x serial dilutions were loaded onto 4–20% TGX^™^ precast mini gels (BioRad, England) either with or without reduction with 50 mM dithiothreitol (DTT). Electrophoresis was carried out at 150 V constant voltages for 60 minutes until the tracking dye reached the bottom of acrylamide gel. The gel was then incubated in sensitization buffer (30% v/v ethanol, 10% acetic acid and 10% methanol) for 1 hour followed by standard Coomassie blue staining procedures. Bands of interest were dissected and collected for sequence identification.

#### Reconstruction of phylogenetic relationships

The mitochondrial genes *cytochrome oxidase subunit 1* (*COI*), *12S rRNA* and *16S rRNA* of *Eoperipatus*. sp. and *Ep*. cf. *barbadensis* were amplified and sequenced as described elsewhere (5). Resulting sequences were deposited in the GenBank (*SI Appendix*, Table S9). Corresponding sequences of *Eu. rowelli* and 34 additional species were obtained from GenBank (*SI Appendix*, Table S9). Primers Translated amino acids of *COI* sequences were verified *a priori* using TranslatorX (35). Amino acid (*COI*) and nucleotide (*12s rRNA, 16s rRNA*) sequences were aligned, concatenated, and analyzed using maximum likelihood inference method as described previously (36).

#### RNA-sequencing, analysis, and sequence identification

The RNA extraction, sequencing, analysis and verification were performed as described previously (37). Briefly, total RNAs extracted from velvet worm glands were reverse-transcribed following manufacturers’ instructions. RNA-sequencing libraries were then prepared and loaded to the HiSeq 2000 followed by data analysis. Sequence identification was assisted with Liquid Chromatography Tandem Mass Spectrometry (LC-MS/MS) as described by Amini et al. with modifications described by Lu et al. previously (19, 37). Sequence verification was performed by cloning and sequencing the respective genes of interest from cDNA obtained using Rapid amplification of cDNA ends (RACE) kit (Takara Bio, Japan) following standard protocol and primers listed in Supplementary Information (*SI Appendix*, Table S8). The naming of different MW proteins has been harmonized as species followed by protein MW region, sequence number and variant number (if any). Sequences previously referred to in *Eu. rowelli* are correlated as following: ER_PH1 (accession: ADI48487.1), ER_PH2 (accession: ADI48489.1).

#### Primary sequence analysis, homology search and structural predictions

Intrinsically disordered protein prediction was performed using IUPred3 webserver (https://iupred.elte.hu/) (38). Sequence fragments with score >0.5 were denotated as disordered regions. Sequence alignment was performed using Clustal Omega tool on the European Molecular Biology Laboratory’s European Bioinformatics Institute website (https://www.ebi.ac.uk/Tools/msa/clustalo/) and graphically enhanced either with ESPript 3.0 (https://espript.ibcp.fr/ESPript/ESPript/) and/or Illustrator for Biological Sequences 2.0 (https://ibs.renlab.org/#/server) (39, 40). Protein sequence logo was created with WebLogo 3 (https://weblogo.berkeley.edu/logo.cgi) (41). Calculation of molecular weights and isoelectric points of encoded proteins were performed using PROTPARAM on the ExPASy server (www.expasy.org). Amino acid hydrophobicity analysis was performed using the hydropathy (Roseman) plot tool on ProtScale (https://web.expasy.org/protscale/) (42). SignalP 6.0 from Technical University of Denmark (https://services.healthtech.dtu.dk/service.php?SignalP-6.0) was used for signal peptide prediction (43). Protein structures were predicted using AlphaFold3. (https://golgi.sandbox.google.com/) (44). In general, structural domains with predicted local distance difference test (pLDDT) scores <50 indicated the low confidence of structural arrangement therefore considered as potentially disordered. For long sequences, the PDB structure files were predicted individually and combined afterwards using ab initio domain assembly (AIDA) tool on the server (https://aida.godziklab.org/) (45). The assembled structures were further colored with Chimera software for display (46). Protein sequence homology search was performed using BLAST function on UniProt (https://www.uniprot.org/) (47). High quality hits were selected by setting the expect threshold (E value) cut-off as <1e -50 with preference given to reviewed entries in the database. The prediction of leucine-rich-repeats (LRRs) was performed using LRRpredictor 1.0 (https://lrrpredictor.biochim.ro/), where hits with LRRpred score >0.5 are considered as high probability predictions and included (48). The LRRs were cross-checked with predicted structural models to exclude regions with low pLDDT score for structural analysis. Protein-protein interaction prediction was performed using PEPPI (Pipeline for the Extraction of Predicted Protein-protein Interactions, https://zhanggroup.org/PEPPI/) (49). Protein functional homology search was performed using COFACTOR (https://zhanggroup.org/COFACTOR/) (50). TM domain prediction was performed using TMHMM 2.0 (https://services.healthtech.dtu.dk/services/TMHMM-2.0/) (51).

#### Circular dichroism (CD)

CD spectra were recorded on an Aviv 420 spectrometer (Aviv Biomedical Inc., New Jersey, USA) with a temperature controller, using a QS quartz cuvette (Hellma GmbH & Co., Mülheim, Germany) with 0.5 mm path length. Two sets of CD conditions were used: i) 30 μM peptide +/-POPC lipid, to test lipid binding and ii) 1 mM peptide incubated on its own, to test self-assembly. Data acquisition was performed in the wavelength range 180–260 nm, with 1 nm steps and averaging time of 1 s. Triplicate scans were averaged and smoothed using the Savitzky–Golay method with a polynomial order of 2. The data are presented as mean residue molar ellipticity (deg*cm^2^/dmol). LRR peptides (LR24: LKKLSLLNNKIKTIRTGTFKDLKR from ER_PM1-var1, LD24: LKDLTLYGNKLKELKQNTFKGLND from ES_PM1, and LN23: LNRLEIDSNNI SEIEQESFVGTN from EB_PM1-var1) were purchased from GL Biochem (Shanghai) and used directly without further purification. POPC (1-palmitoyl-2-oleoyl-sn-glycero-3-phosphocholine) was purchased from Avanti (Alabaster, Alabama) and prepared as described previously (52). Deconvolution was performed using HeliQuest webserver (https://heliquest.ipmc.cnrs.fr/) (53).

## Data availability

The assembled transcriptome and raw RNA-sequencing data is submitted to NCBI under Bioproject PRJNA806368 (ES). The raw spectra and search data were uploaded to the Jpost repository with the following accession numbers: JPST001465 and JPST003160 (jPOST) and PXD031722 (ProteomeXchange). The RNA-sequencing data of *Eu. rowelli* and *Epiperipatus* cf. *barbadensis* is available under Bioproject ID PRJNA157805 and PRJNA1109931, respectively.

## Acknowledgments

This work was supported by the German Research Foundation (grants MA 4147/7-1 and 4147/2 to G.M.), Alexander von Humboldt Foundation (Feodor Lynen Research Fellowship to A.B.), the Natural Sciences and Engineering Research Council of Canada (grant RGPIN-2018-05243 to M.J.H.), a Canada Research Chair award (CRC Tier grant 2 950-231953 to M.J.H.), and the Singapore Energy Research Center (SgEC to A.M). X.G. and R.M.S. thank the support of A*STAR Core funding and the Singapore National Research Foundation under its NRF-SIS “SingMass” scheme (RMS). We thank Gagan Daliaho, Dave M. Rowell, Christine Martin, and Isabelle Schumann for help with specimen collection, Sandra Treffkorn for assistance with RNA extractions and Niklas Metzendorf for cDNA synthesis.

