## Supplementary material for "Receptor-based protein binding in the supramolecular network of velvet worm slime": LRR protein_PNAS template SI_final.docx

**This PDF file includes:**

Supporting text

Figures S1 to S12

Tables S1 to S8

SI References

**Other supporting materials for this manuscript include the following:**

Movies S1

Movies S2

Supporting Information Text

To identify cross-species characteristics of proteins and common essential mechanisms of velvet worm slime, representative specimens for both taxonomical major subgroups have been selected. Maximum likelihood reconstruction of the phylogenetic relationships of selected onychophoran species is based on *COI*, *12S rRNA* and *16S rRNA* mitochondrial genes (Fig. S1, Table S9). For the Australian species *Euperipatoides rowelli* (1), previously published sequences were used. Corresponding sequences of peripatids from Singapore (*Eoperipatus* sp.) and Barbados (*Epiperipatus* cf. *barbadensis*) were additionally sequenced. The resulting tree confirms the phylogenetic position of *Eu. rowelli* within the Australian Peripatopsidae. The species from Barbados is retrieved among neotropical peripatids from Central America and the northern coast of South America. Currently only one species is described for Barbados (2) but since cryptic speciation is a recurrent among peripatids, we cautiously refer to it here as *Epiperipatus* cf. *barbadensis*. The sequenced specimen from Singapore appears within Peripatidae as the sister group to an undescribed representative of *Eoperipatus* sp. from Thailand, which is in line with Lu et al. 2022 (3); we thus also refer to the Singaporean species as *Eoperipatus* sp. The distant phylogenetic relationships of the species studied allows us to gain insights into ~380 MY conserved characteristics and essential functions in the slime common to all velvet worm species.

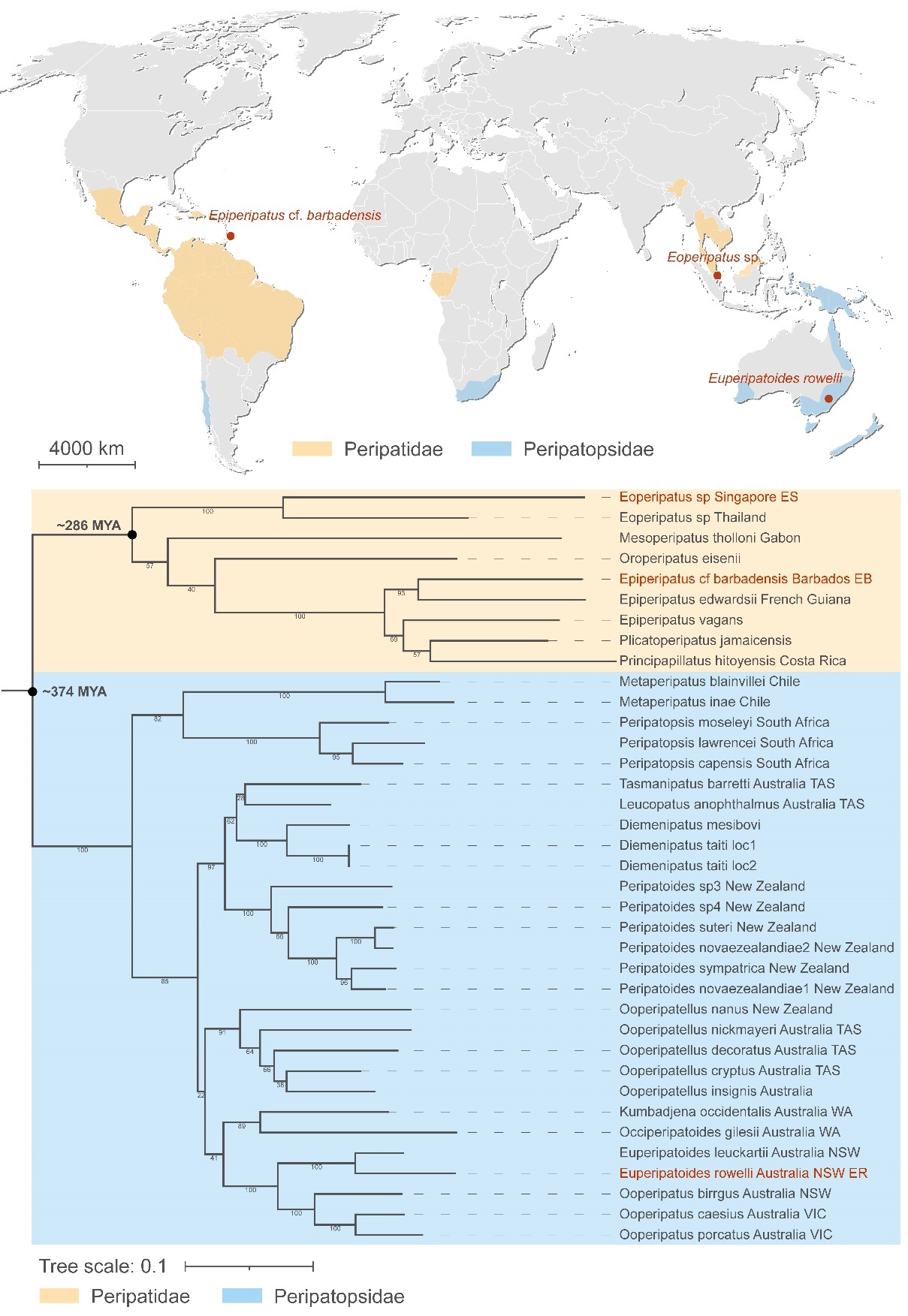

### Fig. S1. Geographic distribution and phylogenetic relationships. Distribution of Peripatidae and Peripatopsidae is highlighted in light orange and blue, respectively. Red dots indicate the collecting sites of specimens used. Maximum likelihood reconstruction of a phylogenetic tree based on amino acid (COI) and nucleotide (12S rRNA and 16S rRNA) sequences of mitochondrial genes from selected representatives of both major onychophoran subgroups. Collection sites of the species studied are highlighted in red. Black dots indicate the divergence date estimated for clades containing the species studied (4).

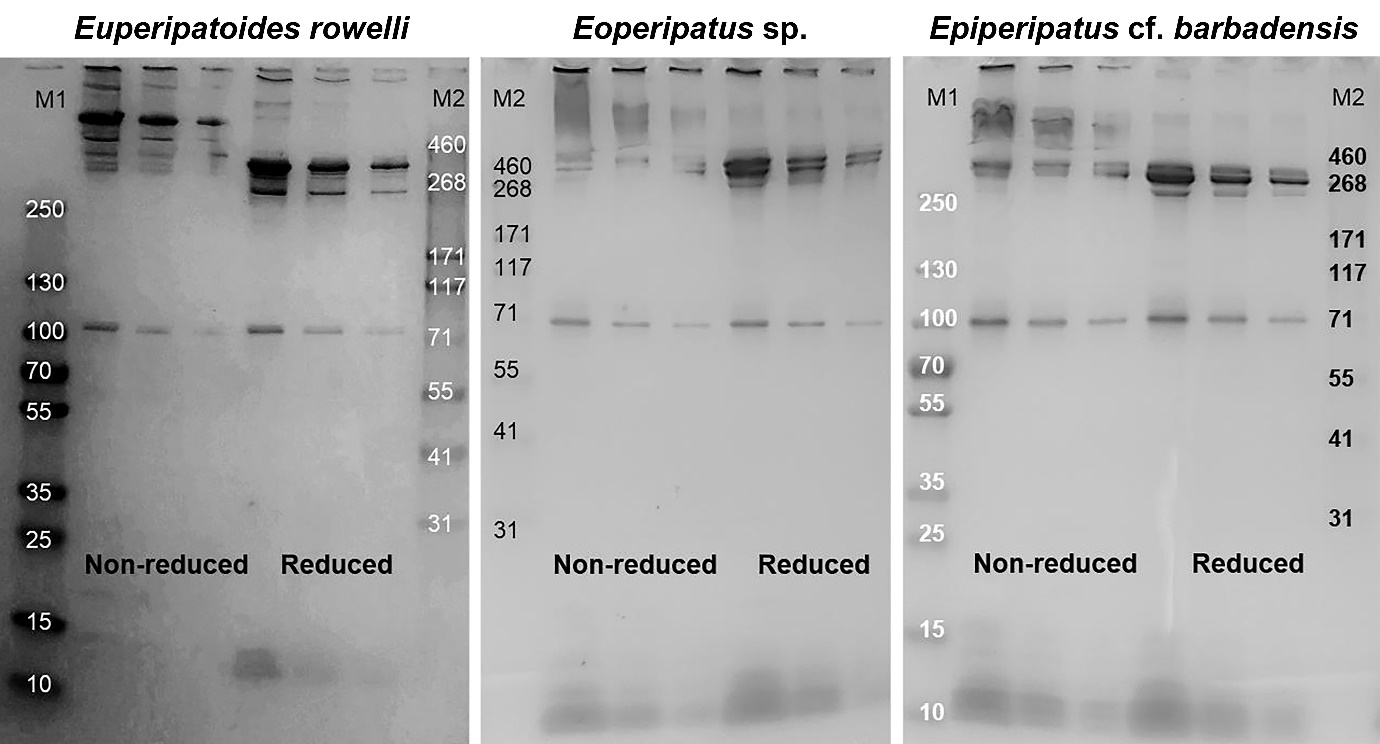

Fig. S2. Full range of molecular weight distribution of slime proteins. Mid- and high-MW are co-conserved across representatives of both major subgroups of velvet worms while low-MW proteins show species specific differences.

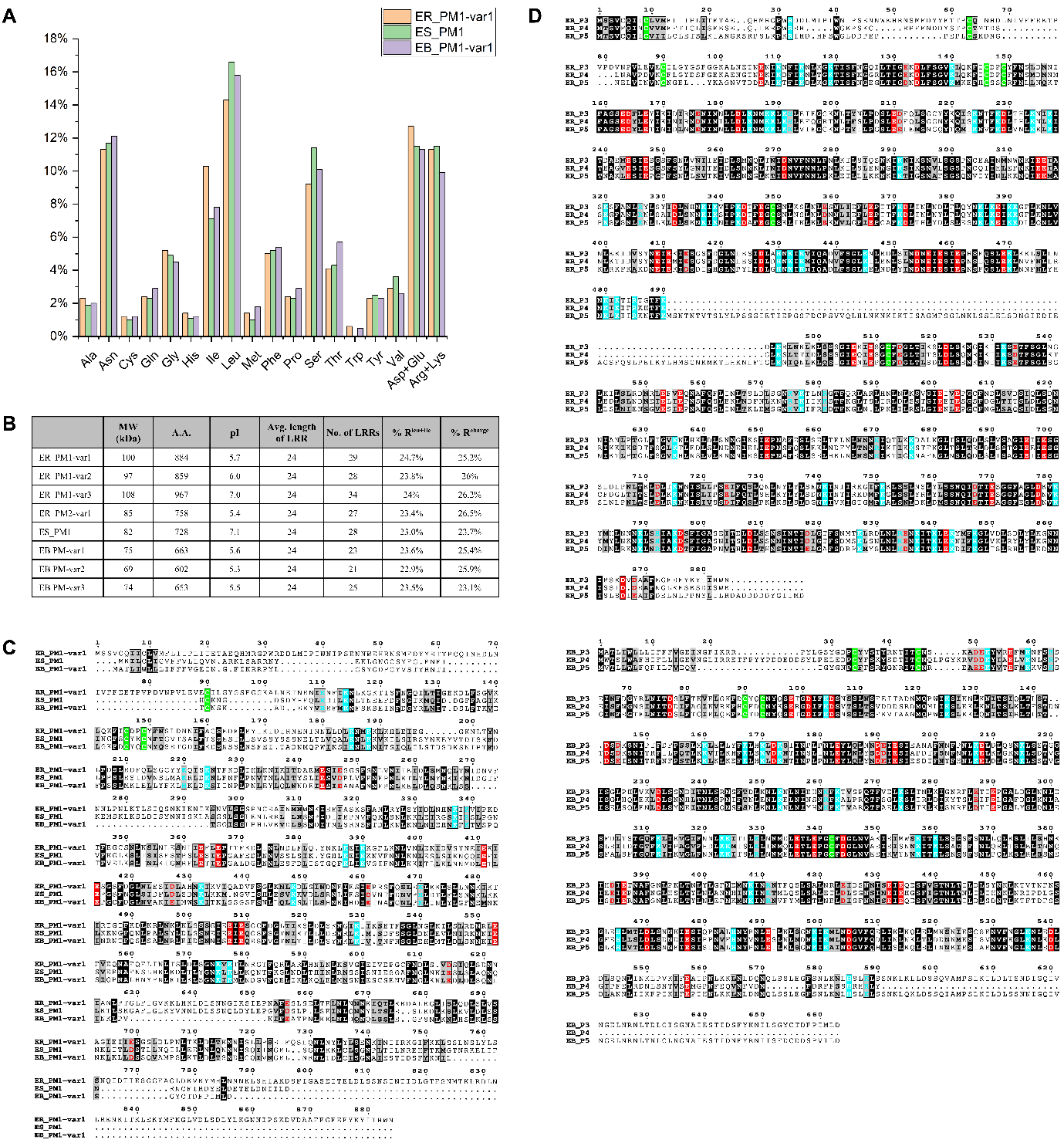

Fig. S3. Sequence analysis of mid-MW proteins. (A) Analysis of the amino acid composition of representative LRR proteins shows conservation across the species studied. (B) Classic parameters of proteins, MW = molecular weight; A.A. = number of amino acid residues; pI = calculated isoelectric point; % R^leu+Ile^ = percentage of leucine and isoleucine; % R^charge^ = percentage of charged residues. (C) Sequence alignment of LRR mid-MW proteins in *Eu. rowelli*, *Eoperipatus* sp. and *Ep.* cf. *barbadensis*. (D) Sequence alignment of potential mid-MW protein isoforms. (Top) ER_PM1-var1–3 in *Eu. rowelli*. (Bottom) EB_PM1-var1–3 in *Ep.* cf. *barbadensis*. Conserved residues are highlighted in colors.

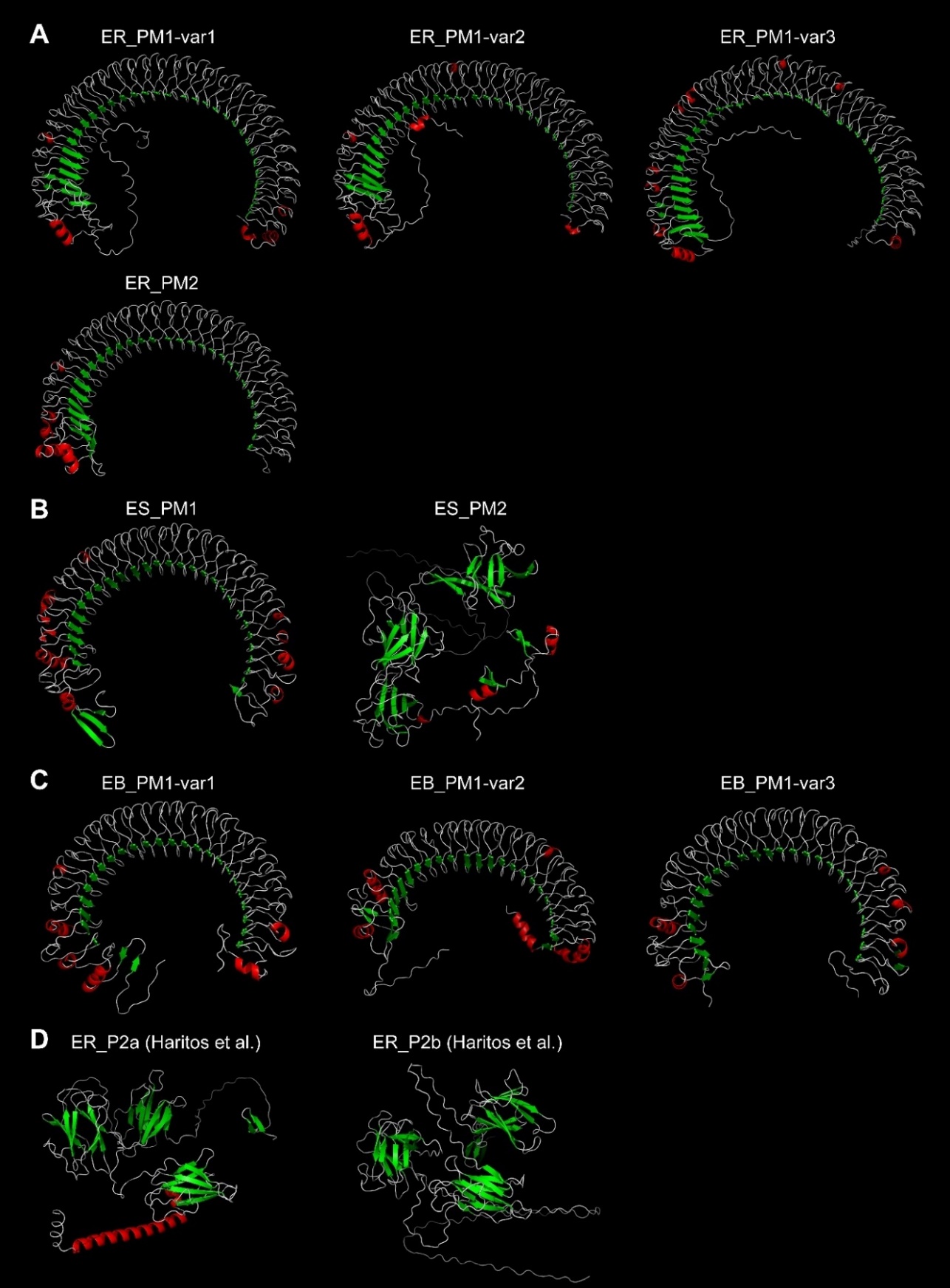

Fig. S4. AlphaFold3 predicted structures of all mid-MW proteins. (A) Left to right: ER_PM1-var1–3 and ER_PM2 in *Eu. rowelli* (B) ES_PM1-2 in *Eoperipatus* sp. (C) EB_PM1-var1–3 in *Ep.* cf. *barbadensis*. (D) Predicted structures of previously reported non-LRR proteins HM217028 and HM217029 in *Eu. rowelli*. Red: β-sheet, green: α-helix.

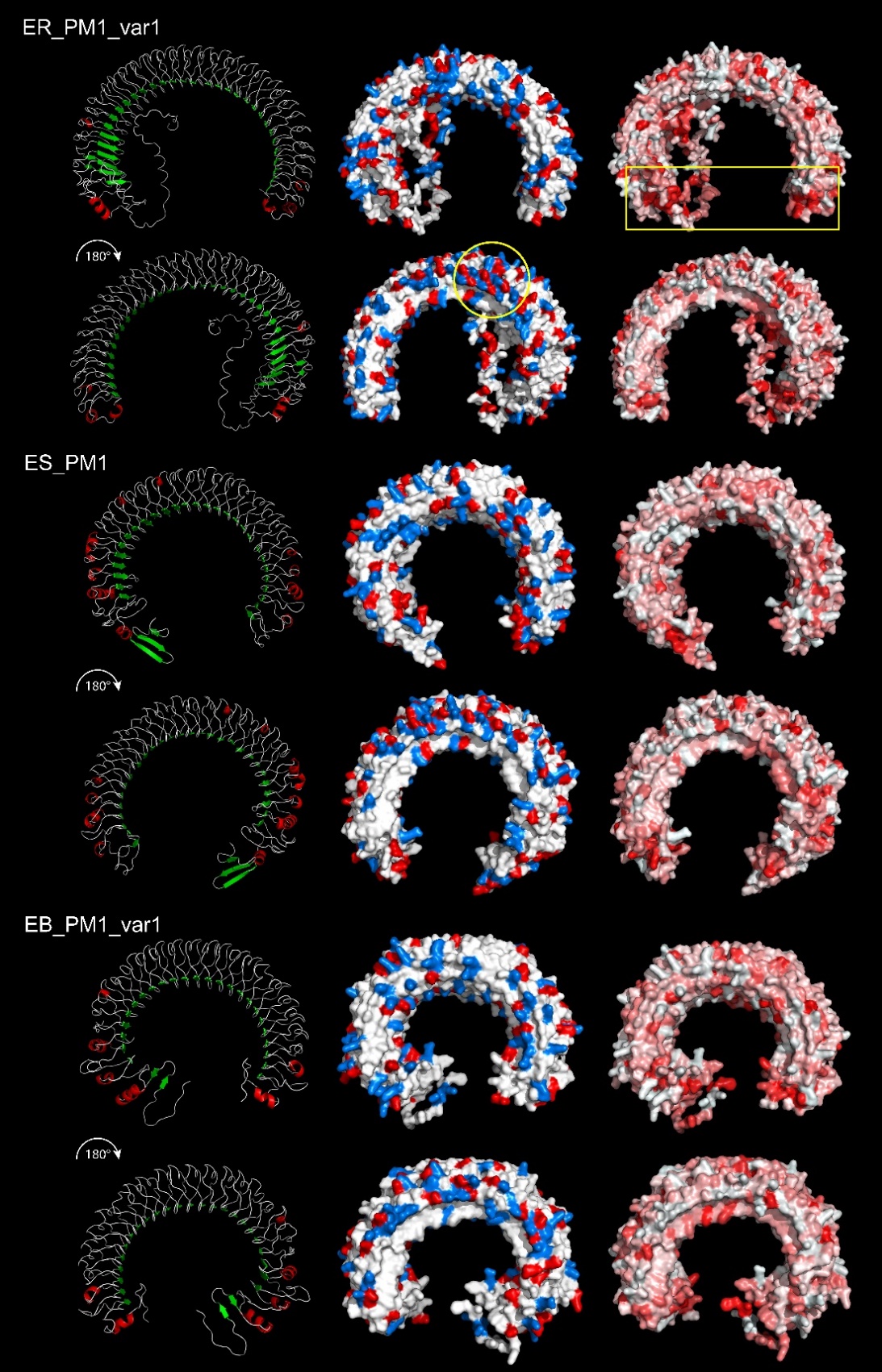

Fig. S5. Structural features of selected LRR proteins in in *Eu. rowelli*, *Eoperipatus* sp. and *Ep.* cf. *barbadensis*. Both sides (turned 180°) of the proteins with annotated secondary structures, α-helices in red, β-sheets in green (left column); Surface charges, positively charged residues in red, negatively in blue (middle column); Surface hydrophobicity, gradient from hydrophilic in white to hydrophobic in red (right column). Yellow rectangle highlights abundant hydrophobic residues at N- and C-termini. Yellow circle highlights a band of residues with alternating charges.

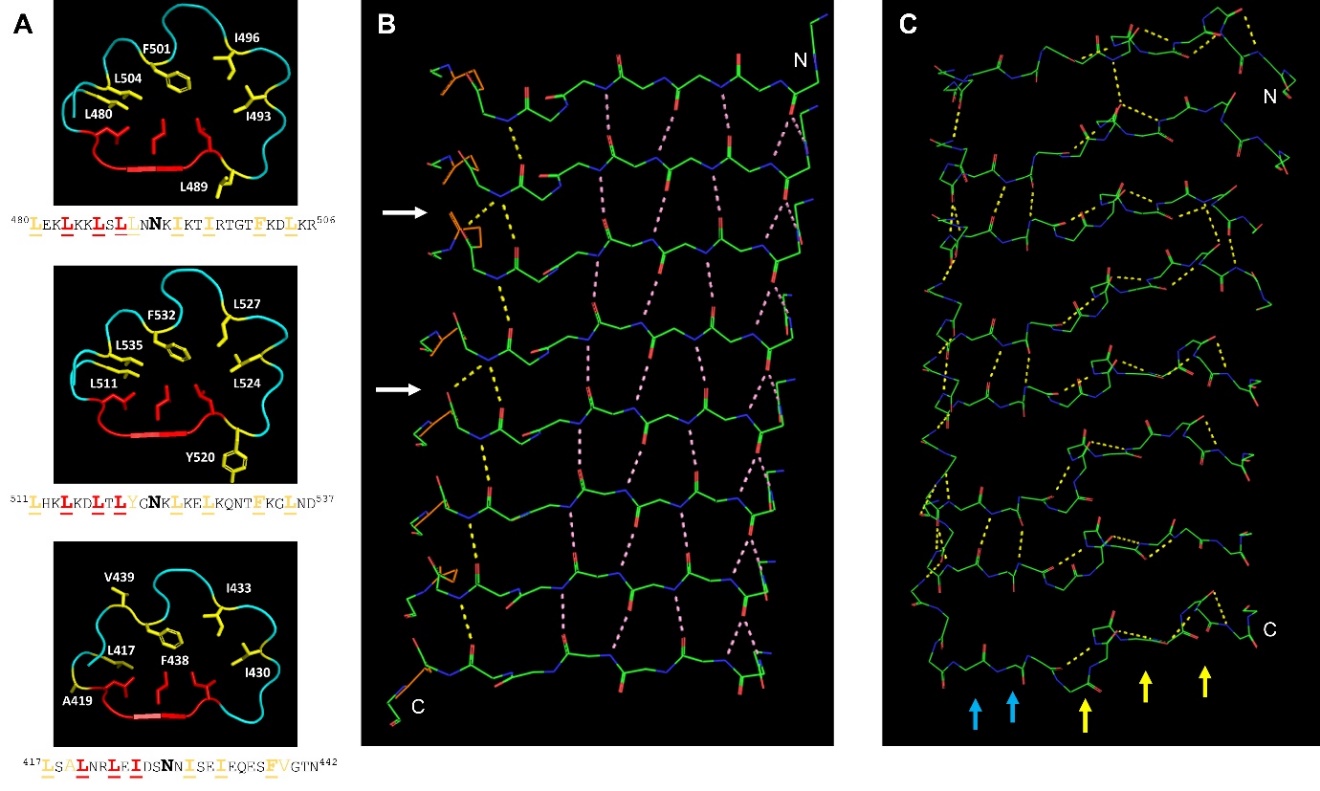

Fig. S6. Structural analysis of LRR loops in mid-MW proteins. (A) Representative loop structure where hydrophobic side chains of amino acids facing inwards. Residues with hydrophobic side chains are in large font size and colored in yellow, and those facing inwards are underlined. Conserved residues are in bold. β-strand structure and corresponding amino acids are labeled in red. (B) Representative main chain hydrogen bonding network between LRR loops in the concave side of ER_PM1-var1. The sequence displayed corresponds to L_554_ to N_728_. Arrows indicate occasions where the fifth C=O accepts hydrogen bonds form two N-H groups. Conserved Asn in orange; hydrogen bonds from β-sheet in pink; hydrogen bonds from either side of each LRR in yellow. (C) Corresponding convex side of L_554_ to N_728_ in ER_PM1-var1. Blue arrows show identified hydrogen bonds that are running perpendicularly to the polypeptide backbone. Yellow arrows indicate positions without hydrogen bonds.

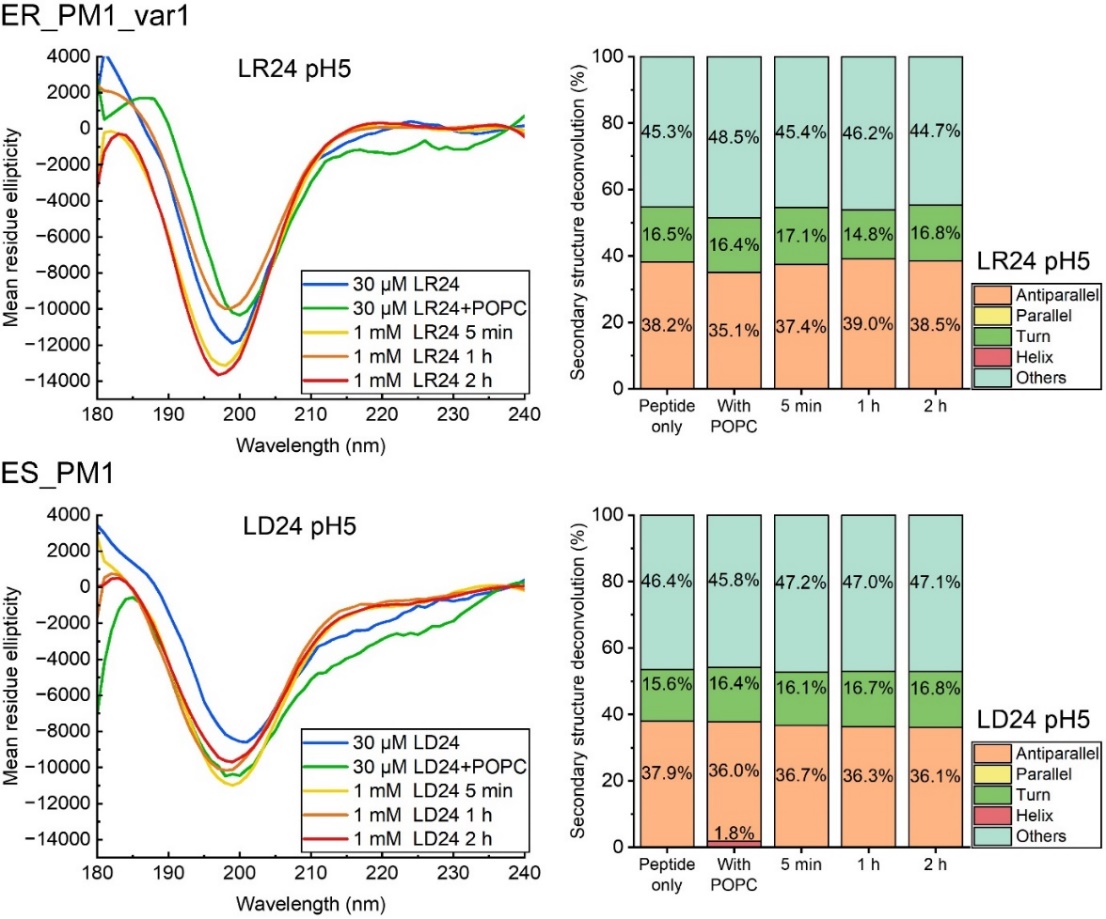

Fig. S7. Circular dichroism (CD) on LRR peptides derived from mid-MW proteins in velvet worm slime. (Above) LR24 derived from ER_PM1-var1 (*Eu. rowelli*) and (Below) LD24 derived from ES_PM1 (*Eoperipatus* sp.) incubated either in the presence or absence of 1-palmitoyl-2-oleoyl-glycero-3-phosphocholine (POPC lipid). The third LRR peptide from *Ep.* cf. *barbadensis* was poorly soluble. No CD data available. Left: CD spectra; Right: estimated secondary structure content (%) after deconvolution.

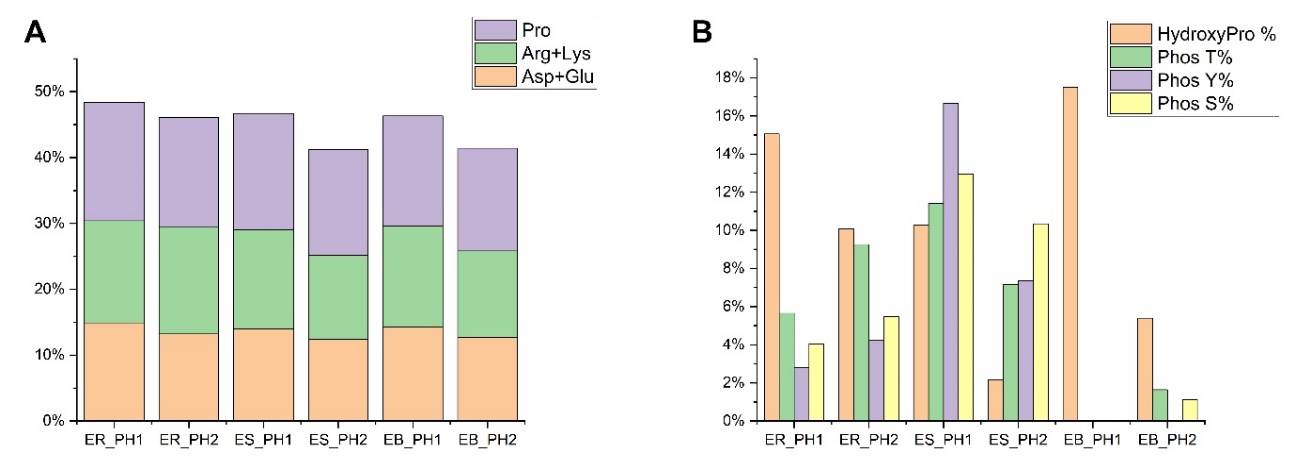

Fig. S8. Amino acid composition analysis of major high-MW proteins in the three species studied. (A) Analysis of charged residual and proline composition. (B) Percentage of hydroxylated prolines and phosphorylated residues, where percentage is calculated as modified over total number of specific amino acid in the sequences. Notably, PH2 sequences show less PTMs than PH1 in all species. Degree of protein phosphorylation is lower for EB_PH2 in relation to the other species and absent in EB_PH1.

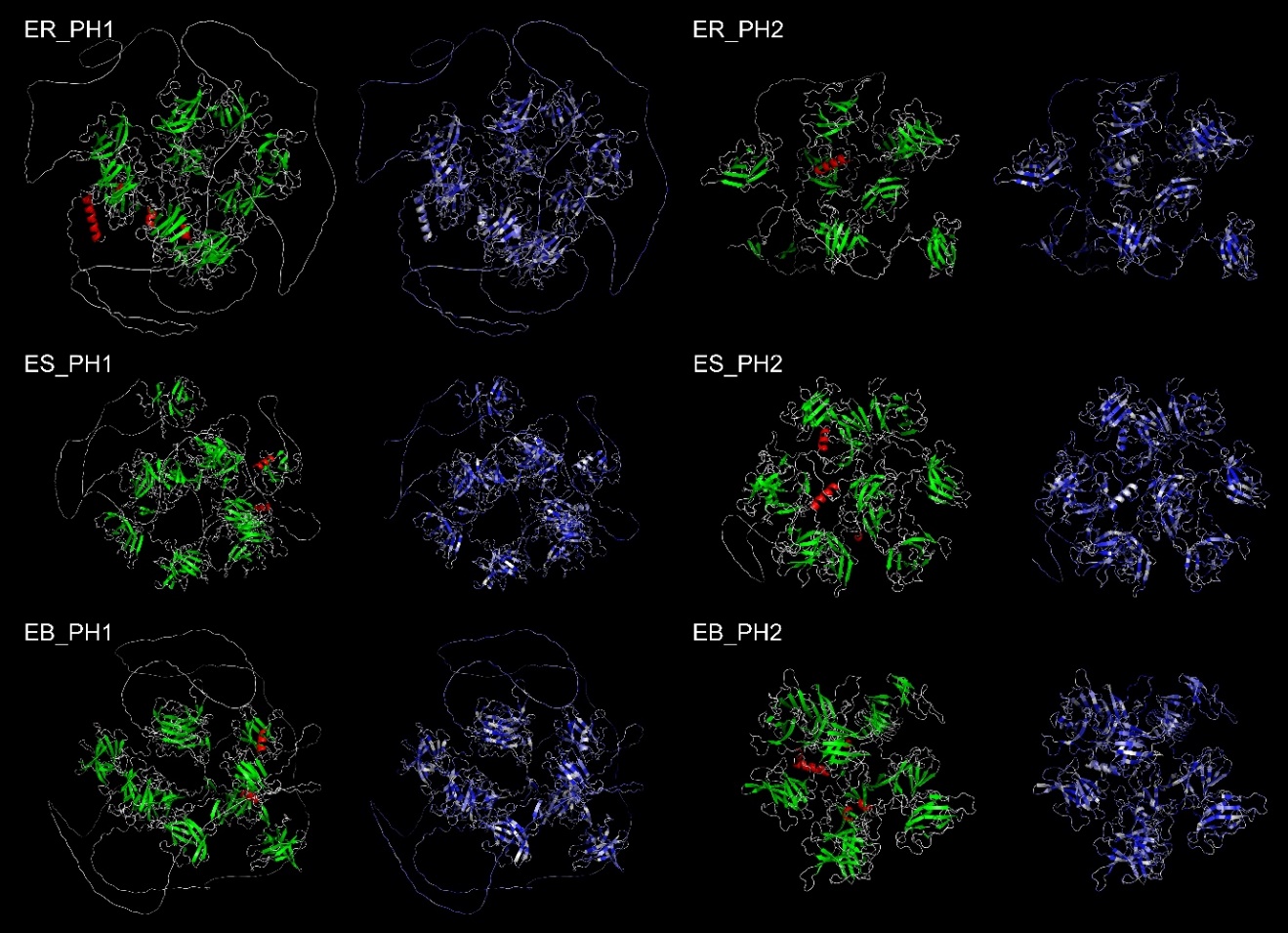

Fig. S9. AlphaFold3 predicted structures of high-MW proteins in *Eu. rowelli*, *Eoperipatus* sp. and *Ep.* cf. *barbadensis*. α-Helices are highlighted in red; β-sheets in green; Hydrophobic residues are highlighted in blue and hydrophilic residues in white

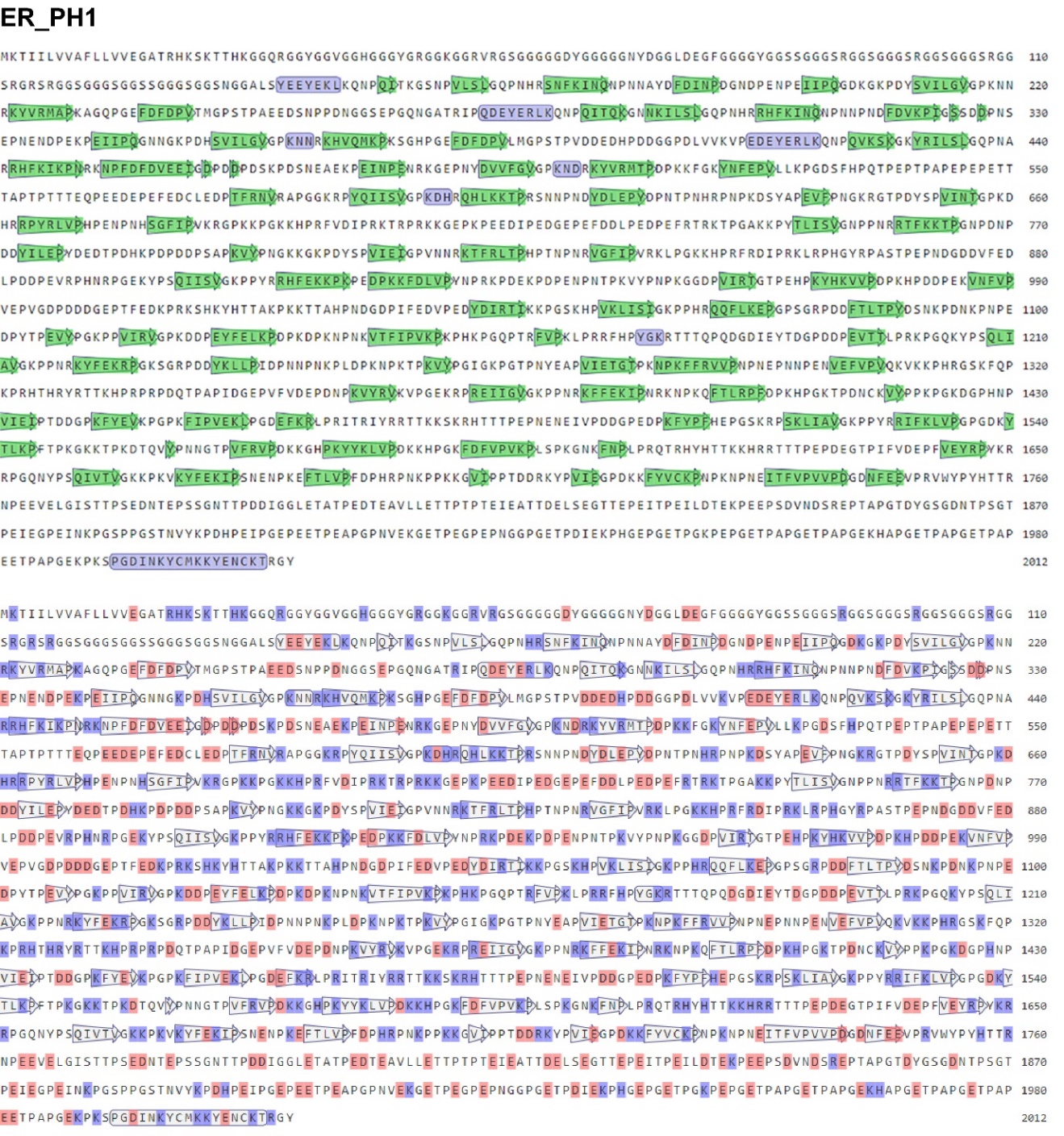

Fig. S10. Annotated secondary protein conformations and charged residues in high-MW protein ER_PH1 from *Eu. rowelli*. (Above) β-sheet in green, alpha helix in purple, (Below) positively charged amino acids in blue, negatively charged amino acids in red; Notably regions of accumulated amino acids of the same charge alternate throughout the core of the protein sequence.

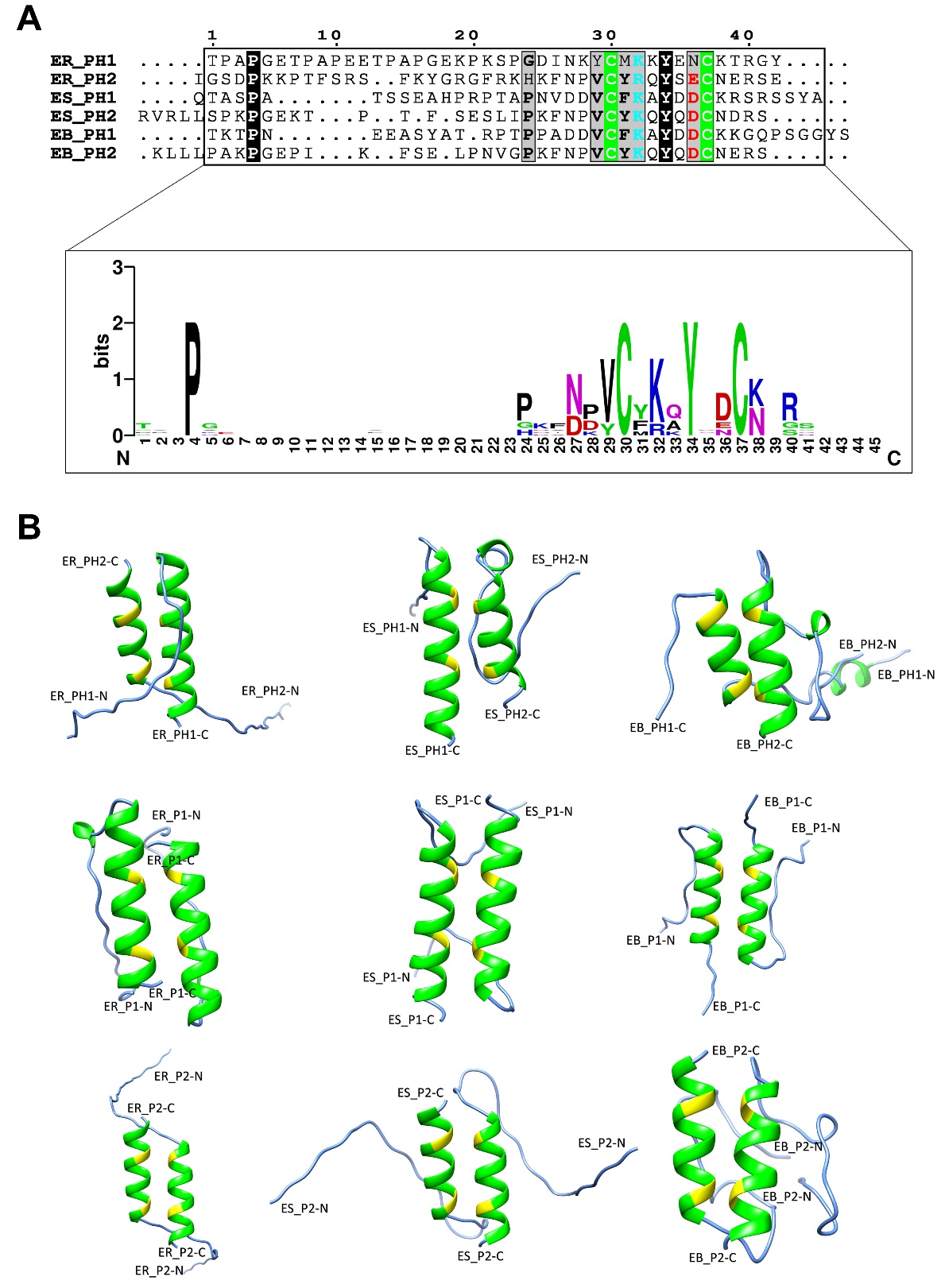

Fig. S11. AlphaFold3 predicted dimerization of C-terminal conserved regions of high-MW slime proteins in the three species studied. (A) The conserved cysteines are highlighted in yellow and random coil in blue. (B) AlphaFold3 predicted examples of possible local structural arrangement of C-terminal sequences in high-MW proteins. Conserved C-terminal motifs (Cys highlighted in yellow, α-helices in green) allow disulfide binding of monomers into protein complexes.

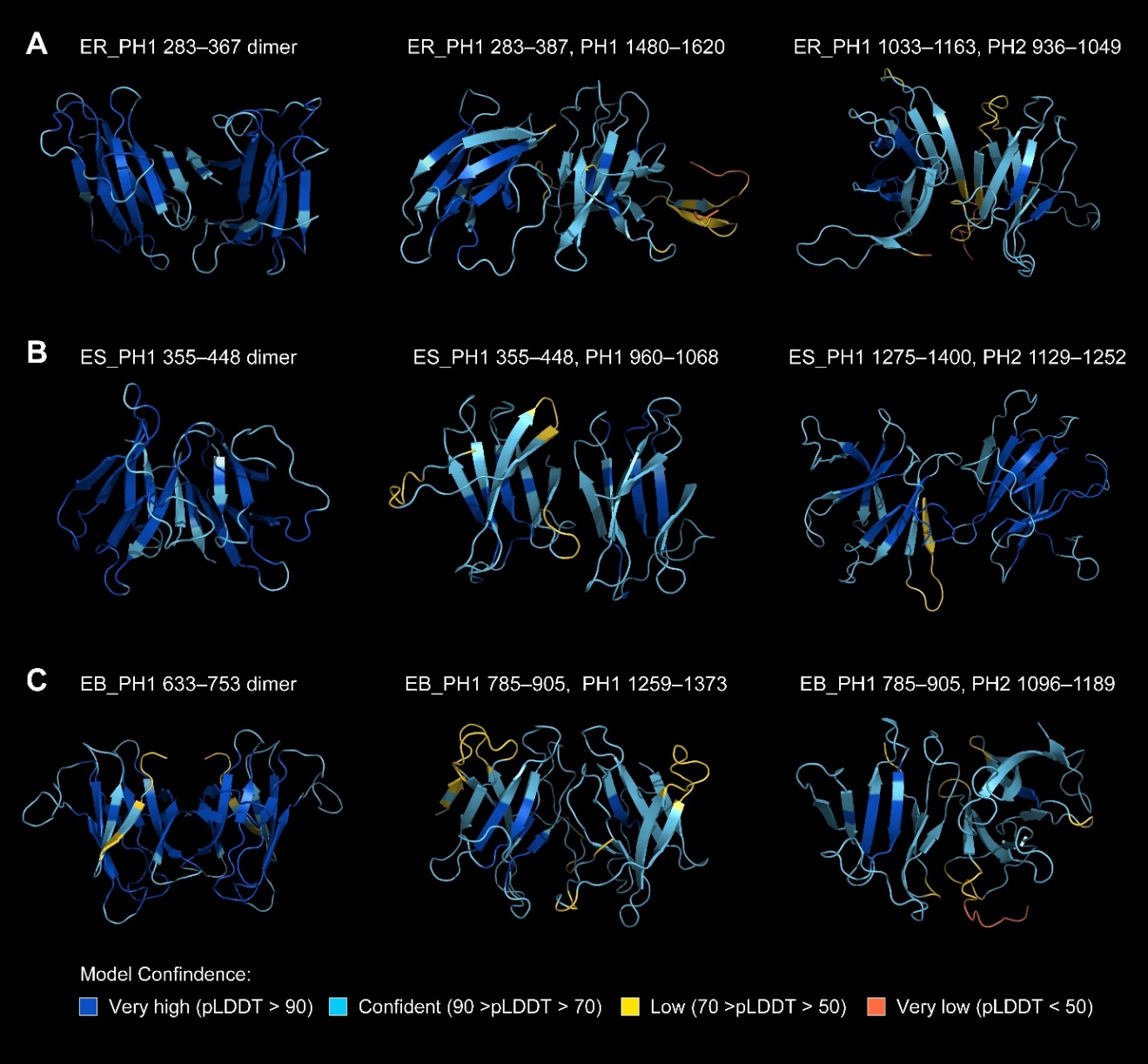

Fig. S12. AlphaFold3 predicted β-sheet domain stacking is prevalent in the high-MW interaction network. Representative structures of respective species, (A) *Eu. rowelli*, (B) *Eoperipatus* sp. and (C) *Ep.* cf. *barbadensis*. Left to right: β-sheet domain homodimer, heterodimer in the same and different high-MW proteins. Sequences are displayed on top of each figure. Dominant blue coloring indicates high confidence prediction with AlphaFold3.

Table S1. Mid-MW protein sequences

**>ER_PM1-var1 (accession: PP712084.1)**

MSSVCQIICLVMFLIIPLITETAEQHMRGPWRDDLMIPIWNIPSENNWEHRNSMFDYYETTPCQINHDLNIVFFEETPVPDVNPVLEVECILGYGSFGGKALNETNENIKNFIKNLKGKTISFNGQILTIGEKDLFSGVKLQKFICDPCYFNSLDNNIFAGSEDFLEYIKIDIRNENINNLLDLKNMKKLKHLEIEGGKNLTYNLPDSLEDFQLSGCYYKQISKNTFKDLTHLKNIKITDAEMESIESGSFSNLVNIIEIDLSMNQLTNIDNVFNNLPNLKTLSIQSNKIKNIKSNVLSGSPNCEAINMNWNKIEEIASKSFANLRYLSYIDLNHNKIKVIPKDTFEGCSNLKSLNLESNLIEFLEPTTFKDLINLNDLFLQYNKLKEIKKGFLKNLVNLEKIDVSYNEIEKIESGSFDGLNLESIDLAHNKIKVIQADVFSGLKNLKDLSINDNEIESIEPRSFQSLEKLKKLSLLNNKIKTIRTGTFKDLKRLNKLKLSSSGIEEIESGCFDGLTIKSLDLSKNGIKIIKSETFSGLNGLKILSLRDNRLEFVEQNAFQFLINLTSLDLSGNKVKTLNRGTFQRLARLHNLNLKSVGIEIVEPGCFNDLSVDSIQLSDNNIANLPTGLFIGVKKLHKLDLSNNGIKSIEPNAFESLSELTELNLNNNKIQTLKKDALKGLISLQDLSLVSAGIEIIESGSLDLPNLTKLDLTKNNISLLPSEIFQSLQNLNYLYLSENKINIIRKGIFKKLSSLNSLYLSSNQIDTIESGGFAGLDKVKYMHLNNNKLSHIAKDSFIGASEITELDLSSNSINTIDLGTFSNMTKLRDLNLRENKITKLEKYMFKGLVDLSDLYLKGNNIPSKDVDAAFEGFEFYKYIIHWN*

**>ER_PM1-var2 (accession: PP712085.1)**

MTSVFQINCVVMFFTIGLISEESEQLKRPWREHWGKPSKGRPFFNDDYSTPCTMTDSLNAVPDVKCFLGYDSFGEKAENGTNEKIKDFIKNLTGKTISFKGGRLTIGEKDLFSGVKLQKFICDPCFFNSMDNNMFAGSEDYLEYIKINIRNDNINILLDLKNMKKLKHLEFQGRTNLTFSLPDSLEDFQLSGGNYKQISKNTFKDLIHLKNLKITEAGVESIESGSFSYLGNITEIDLSNNKLTNIDNVFNNLPNLKILSLENNGIKSIKSNVLSGSPNCQIIHLEFNKIEEIASKGFANLRNLSAIDLSNNKIKSIPKDTFEGCSNLNSLNLDNNLIEFLEPTTFKDLINLNYLFLQYNKLKEIKKGFLKNLVNLKTIDVSYNEIEMIESGSFDGLNLESIDLDHNKIKMIQANVFSGLKILENLSLNDNEIESIEPNSLQSLEKLNVFNLLRNKIKIFRKHTFKGLKSLTTIDLSSSGIEQIESGCFEGLTITSLDLSQNRIKIIKSDTFSGLKTLEDLSLNDNEIELIEPNSFQSLEKLNDFNLKNNRIKIIKKSTFKGLTSLTRLDLSSTGIEEIESGSFDGLTITSLDLSQNRIKKIKSDTFSGLKTLEDLSLNDNEIELIEPNSFQSLEKLNDFYLTNNRIKIIKKSTFKGLTSLTRLDLSSTGIEEIESGCFDGLTITSLDLKKNNISLIPSELFQTLSHLKYLYLSDNKINTIRKDMFKGLSSLYRLYLSSNQIDTIESGGFAGLDNVKYMYLNKNKLSHIAKDSFIGASNITSLDLSSNNINTIDIGTFSDMSKLSNLNLKENKITKLEKNIFKGLVKLRYLYLKGNNISSIDDEAFNGLKFSKSDISWK*

**>ER_PM1-var3 (accession: PP712086.1)**

MTSVCQIICVIILCLETSLKIANGRSRPSLRPKRIHDHFSWGLDDPEPPEPLCSKDNGNELVINVNCNGELYKAGNVTDDEAIKTFIKDLKGKTISFNGEGLTIGDNDLFSGVKMKEFHCSSCRFNLLNQKTFAGSEDYLETINIDLNNENINNLLDLKNMKKLKKLVIEGGKNPTYILPGSLEELEMSNGQYTQMIKNVFKDLVNLKKLKITNAKLESIEPGFFSNLLSVTKIVLSNNGLKTIDNVFNNLPKLETLLLKKNKIKTIGSNAFSGSQNVIYINLKKNQIEEMAPKSFSNLKNLLKLDLSKNKIKNIPKDAFEGCSKLIKLKLEKNVLGFIEPGAFKDLTHLYDLSLESNKLKEIKKDLLQNLVKLRKFKAKDNEIEKIDSDIFHGLNPYLIDLGHNKIKIIQADLFSGLKELNDLYINDNEIESIEPNSFQSLEKLNNFNLYHNKLKIIRKNTFKMSNTNTVTSLYLPSSGIETIEPGSFDCPSVVQLHLNKNKIKTISAGMFSGLNKLSSLELSDNGIEDIEAGSFQSLPELKYLHMSGNKMKTLHKNLFTGLKNIQNLKLQSIGIENLEPGCFDGLTLSDLDLSENKINTIKSETFSGLSGLLSLNIENSGVESIEPNAFQSLINLTKLEMSGNKVKIFRKDTFKGLNLPKLILRSLGIENLEPGCFNGLSAQSIDLSSNNIKTLPTGLFSGVKHLNSLVLKNNNIESIEPNAFESLSELHKLNLNSNKIKTIGKNAFNGLNSLQDLSLISAGIEIIESGSLNLPNLTSLELRKNKISIVSSDIFQSLPKLKSLSLDGNKIVKIGTGMFKALSNLDKLDMSSNQIETIEAGGFEGLDNVKDINLRKNKLNRIAKDTFIGAPNVTHLDLTHNSINTIELGAFSDRPKMYSLNLKDNKITKLEKDIFKGLTSLRHLTLEDNNISLSDIEAIFDELNLPPNYLTLRDADDDDDYGIIMD*

**>ER_PM2 (accession: PP712087.1)**

MQCWASTWWFKLLILALFSSYVLANVSCPEVESSPCVCTSKNPSGLDIKCHACTNVNDFSAEEKYAKAQKEEAYLKMINGTINSLLSHCILLEEGFFSDAKIKRFHCVDCDLSASSTSVYAPLENILEELKLDDCTIPSIHPFQNLKTLKSLKIDSDNLLVSNEKIVIPSPLEHFAIYNAKFNQVFKDLFTTYSSLTTLKIQSTNLQKLEPNCFQSLKNLTLIDLSMNRLTKIDDVFKNLPELQELFLEQNRIKTLSPQTFTNMKKLKFIDLDMNEIEKIEAKTFSNLPNIVSIALGENNIKEISKDAFLGADEIKELNLGNNEIESITPSPFAKLAKLERLNLAVNKIKSITSETFSDCLNLMDLNLKSNNLQYLEPKMFSHLKKLKRIDLWHNGIKEIKKDALDGLLKLTSFSLVSNSLELIDPGAFEGLESVEILDLHDNKMTNINKGIFSDMPKLHTLLINNNKIEKIEPGAFKDLPNIDHLNLNFNAMKKLETETFVGASMLKELDLEENPIEVIEENAFHGLESIVKISLSKAKLSELTNKILSGLPSLTTFRCFSSDLKTIEPGAFKSTNELHDLYLRYNKITKIRRFNFKGLSSLEDLDLAYNNIVRIEPDAFMGLSVLHKLDLSWNKITQTNKDIFLGLTNLKILQLDNNEIQILQPDTFNGLESLETLDLSHNRIADINKEVFEFSNLQVLELESNLLWKVEDDAFDKAPTLTSLNLHGNGINEIAPTAFKNLVNLDKYHVLTSKTSL*

**>ES_PM1 (accession: PP712088.1)**

MKILCLIGVFFVLLQVNARKLSARRNYEKLGNGCSYFGENEIHCKNGDSDYHFQLKDILKNLTNEEFDFSGTKMQIDDGFFAGIKINGFSCYKCNFTGVTSALFTSSESTLLSVSSYSSSNELTLVQALKNLKKVKTLSIRSYNKEYVTDISKFDLPPSLESIDVNSLMAKRLDKNLFNFLPNVTNLAITYSLIESVDPLVFENFPNLKTLYLNYNKIKNIDGTFKEMSKLKSLDISYNNISKLASGSLSGLENLRRLILNSNPIDIIEPNVFQKLSNLKKLEIRGSKIRSLPGNVFKGLKSLTSISFSTPSLESIEPGAFEDLNNVSSLSIKYGHLKEIKKNVFNNLKNLESLSIKNQQIEFIEPGSFNGLSIDFLDLSDNKLKKMNGVISHLESVKKVDLSRNLIESIDVNTFDNVPHLDTLFLSNNKLKKLKKNGFQNLPALSYLDLSSNEIEEIEQGSFSGTNIKTLLLDENRLKMSDFSSGLDNLEVLNIRKNNIESVEPNAFNSLHKLKDLTLYGNKLKELKQNTFKGLNDLTHINLRNSSISNIESGAFNGLNKIESLSLAQNQLKTLSKGAFLGLKYVNNLDLSSNQLDYLEPGVFDSLPLSEINLQNNYLTSLESLKDLKSLKVLDLSSNKLTTLDSTLLNQILLEELNLQNNNISQIINGELSGNLKKLCLSGNPLTILNREHFTKMGTNRKELIFNRNCFIRDYELDETELDNIILD*

**>ES_PM2 (accession: PP712089.1)**

MTMKAFILIVVLCGIAISNAESDPTPEELLAQGYVLQPDGSYLKSGESETVQEGGENMSEDELKAQGYQLQPDGTWLKTETSTETQTQTVRETQKVDYDDNEYQSTNFNNGILTVQSGSDKGYFKINKGASNNENDYSVQAVNSPSDPTPISNPDANSPQVYPQGNQGDPDYSVVLGVGKGKTRNYLRMQPDEARPSGYKFIPQYISGYRPGEATNAPEPVEEPAPGPSDDVEEDEDDNPDEPVVSTNRGKLILSVGRPNRRYFKITPSNNNNPYDFGVEPIKSPSEPDAKPDPNDDEAPEVIPNGNRGSPDYQPILAVGPKDNRRLLGVVVNPKNPRKFKIITREQEEQPGEVQTDPPEEDIDYDDEPPNVNYKVLNIAGKKNEIVSAGPLGKRFHLRIIHRSNRPGDVEIRPIHSAENIDSPPDSNDLHAPEVVRQGNPEDPQNAPIIASGPRYRRVYHKMLIIRGRVVFKPLRRVRIKGRYRFLLRRRRWHWKLGHRRVLRRTKMIIKKRTNNCWFFCGGGGWGWGQQKRKVIVRRPAMMVGSGGANVHLQNRRVVLKTTTQTKPSGKFLFGFFG*

**>EB_PM1-var1 (accession: PP712090.1)**

MATLIWLLLIFFFVGEINGFIKRRPYLGSYGDPCYVSTYRNTITCNSKADEKYVREFMKNFSKSEINFDGYRLNITDSLFTKVELGKFDCYQCNYQSETGDIFKDSNSSLNSFEITADNMQPWIKSIKNLKNITSIQLTLSTDSDKSNIPTFDFPSSLMKLELLYFKLHKLDKSIINPLPNLEYLQLNNDEIESIEANAFNNFPNLKELDLQSNKLSSTGGISGLPHLVKVDLSSNDITNLSRNSFTDLKNLKNLNIEHNKFKTVSPQTFVELKSLTNLKIGNRFLETIEPGALDGLNNLESFDLTSTGQFKLIKVGIFNNLKKITLLKLNNMDLETLEPGCFDGLNVAKIEIMWSKITKLSSGSFSNLLQLKSLILSHNKIHDIETNAFGNLPKLTNLYLGFNEMNKINRNTFQSLSALNRLEIDSNNISEIEQESFVGTNLTILDLSYNKLKTVTNFNSGLEHLMTLDLSNNKIESIQPNALHNYPNLEILKLSGNKIKMLNDGVFQELIKLKQLELMNSNIESCSENVFNGLKNLEDLDLSQNLINKLPVKIFEATPNLKKLNLDQNQLSSLEGFSNLKNLHSLKLSSNKLKLLDSSQVAMPSLKTLDLTSNDIGQIVNGELNRNLTDLCISGNAIESTIDSFYKNILSGYCTDFPIMLD*

**>EB_PM1-var2 (accession: PP712091.1)**

MWTSLWAFHIFLLIGEVNGLIRRETFPYYPDDEDDSYLPELEDPCYFSKYSDTITCNQLPGYKRVDDKYIAELVKNLSRPEISFWGNSINITDDIFAGIKVRKFHCFDCNYKSERGDIFKDSVTSLTSVDDDSRDMQMLIKSLEKLKNLTSLKLTFSTDWTDSDKSNITRFILPQTLKKVTLKHFKLSKVDKNTINSLTNLEYLNLYSDDIESIAPDTFSNFNNLKELKLGKNKLSDTVGISGLHQLEKLDLSNNHLTNLSPNIFTNLENLKELNINSNRFKALPRQTFSGLKSLIRLDIKYTSLEIFELGAFDGLKNLASLEIIDTGTFKVIPAGVFHDLKKMISLKLINMQLETLEPGCFDGLNVAEIEISNNKITKLSAGIFLNLQRLTSLELTNNQIEEIEPNAFNGLHSLTYLNLANNHINKINKTMLQSLSALETLDLSGNSINEIEHGSFIGTNLTNLILIHNKLNRIPDLGSGFEHLLSLDLSDNDIESIPPNVFHNYVNLESLNLNDNKIKMLNDGVFQDLINLNKLTLDDNNMKSLSVNVFKGLKNLKDLGIPFELRDNLSNTVSEMGPNFEQVNFVDNFDRFFSSHRHRLY*

**>EB_PM1-var3 (accession: PP712092.1)**

MVTLLWLFQFILFVEQVTGYGNPCFFSRYGNRITCNREAEEKYVTEFMKKVSKSGINFEGTELNITDSLFTGIELQEFECTDCNYQSETGDIFKDSNSSLTSFKVTAANMQPWIKSLKKLQNITSIRLTLTTDSEISNITRFNFPSTLKKLKLKEFKLHKLDKNTINPLPNLEYLQLYNDEIESIESDTFNYFHNLKELDLGSNKLSSTVGISELSQLENLDLSYNEITNLSQNSFSNLKNLKNLNIYHNKFKTVARQTFAELKSLTKLKISNRFLDTIEPGALYGLDNLESFALSFTGQFKIIKVGLFNNLKKITSLRLNNMELETLEPGCFDGLNVEEIKVTNNKITKLSNGSFSNLPQLKSLRLSSNEISDIERNAFGNLLKLTYLNLEFNKMNKINRNVFYMLSTLNELDISFNNISEIEQESFVGTNLTILDLSDNTLKTFTDFSSGLKHLVTLDLSNNEIESISLNALHNYPNLEILKLSGNKIKAINDGVFQHLISLKQLELINNKIESLAENVFNGLKNLEDLDLANNLIIKFPIKIFEPTHNLKKLNLDNNQLSSLEGFSNLKNLHSLKLSSNKLQKLDSSQIAMPSLKTLDLSSNNIGQIVNGELNRNLTNLCLNGNAIESTIDNFYRNIISFDCDDSPVILD*

**Annotations**

P: hydroxyproline	T/Y/S: phosphorylated residue

C: cysteine	Predicted signal peptide

Underlined: glycine/serine (GS)-rich domain

*Stop codon

Table S2. Predicted amino acid composition of mid-MW proteins.

| **Residue** | **ER_PM1-var1** | **ER_PM1-var2** | **ER_PM1-var3** | **ER_PM2** | **ES_PM1** | **ES_PM2** | **EB_PM1-var1** | **EB_PM1-var2** | **EB_PM1-var3** |
| --- | --- | --- | --- | --- | --- | --- | --- | --- | --- |
| Ala | 2.30% | 1.40% | 2.60% | 4.60% | 1.90% | 4.00% | 2.00% | 2.20% | 2.10% |
| Arg | 1.80% | 2.10% | 1.60% | 1.50% | 2.30% | 7.60% | 1.50% | 2.80% | 2.00% |
| Asn | 11.30% | 10.50% | 11.00% | 9.00% | 11.70% | 7.30% | 12.10% | 10.80% | 12.60% |
| Asp | 5.70% | 6.20% | 5.80% | 5.80% | 5.20% | 6.40% | 5.40% | 7.50% | 4.60% |
| Cys | 1.20% | 1.00% | 1.10% | 1.70% | 1.00% | 0.50% | 1.20% | 0.80% | 1.10% |
| Gln | 2.40% | 2.00% | 1.90% | 2.10% | 2.30% | 4.70% | 2.90% | 1.80% | 2.90% |
| Glu | 7.00% | 6.60% | 6.50% | 7.80% | 6.30% | 6.70% | 5.90% | 5.80% | 7.20% |
| Gly | 5.20% | 5.70% | 6.20% | 2.90% | 4.90% | 8.70% | 4.50% | 4.50% | 4.00% |
| His | 1.40% | 0.90% | 1.60% | 2.10% | 1.10% | 1.40% | 1.20% | 2.30% | 1.20% |
| Ile | 10.30% | 10.00% | 9.20% | 7.80% | 7.10% | 5.50% | 7.80% | 7.80% | 7.80% |
| Leu | 14.30% | 13.70% | 15.00% | 15.60% | 16.60% | 5.20% | 15.80% | 15.10% | 15.80% |
| Lys | 9.50% | 10.10% | 10.80% | 9.40% | 9.20% | 6.10% | 8.40% | 7.50% | 8.10% |
| Met | 1.40% | 1.30% | 1.60% | 1.60% | 1.00% | 1.40% | 1.80% | 1.50% | 1.10% |
| Phe | 5.00% | 5.90% | 4.30% | 5.30% | 5.20% | 2.80% | 5.40% | 5.80% | 6.10% |
| Pro | 2.40% | 1.70% | 3.20% | 3.40% | 2.30% | 9.50% | 2.90% | 3.00% | 2.00% |
| Ser | 9.20% | 10.90% | 9.10% | 8.40% | 11.40% | 5.90% | 10.10% | 8.60% | 9.80% |
| Thr | 4.10% | 5.60% | 4.40% | 5.50% | 4.30% | 4.50% | 5.70% | 5.80% | 6.40% |
| Trp | 0.60% | 0.30% | 0.10% | 0.80% | 0.00% | 1.00% | 0.50% | 0.70% | 0.30% |
| Tyr | 2.30% | 2.10% | 1.40% | 1.50% | 2.50% | 3.60% | 2.30% | 2.30% | 2.10% |
| Val | 2.90% | 1.70% | 2.70% | 3.30% | 3.60% | 7.30% | 2.60% | 3.30% | 2.80% |

Table S3. Summary of LRR motifs identified in mid-MW proteins.

| **Protein** | **L0 position** | **-5** | **-4** | **-3** | **-2** | **-1** | **L** | **x** | **x** | **L** | **x** | **L** | **+6** | **+7** | **+8** | **+9** | **+10** |
| --- | --- | --- | --- | --- | --- | --- | --- | --- | --- | --- | --- | --- | --- | --- | --- | --- | --- |
| ER_PM1-var1 | 179 | E | D | F | L | E | Y | I | K | I | D | I | R | N | E | N | I |
|  | 201 | K | N | M | K | K | L | K | H | L | E | I | E | G | G | K | N |
|  | 220 | N | L | P | D | S | L | E | D | F | Q | L | S | G | C | Y | Y |
|  | 244 | K | D | L | T | H | L | K | N | I | K | I | T | D | A | E | M |
|  | 268 | S | N | L | V | N | I | I | E | I | D | L | S | M | N | Q | L |
|  | 291 | N | N | L | P | N | L | K | T | L | S | I | Q | S | N | K | I |
|  | 315 | S | G | S | P | N | C | E | A | I | N | M | N | W | N | K | I |
|  | 339 | A | N | L | R | Y | L | S | Y | I | D | L | N | H | N | K | I |
|  | 363 | E | G | C | S | N | L | K | S | L | N | L | E | S | N | L | I |
|  | 387 | K | D | L | I | N | L | N | D | L | F | L | Q | Y | N | K | L |
|  | 411 | K | N | L | V | N | L | E | K | I | D | V | S | Y | N | E | I |
|  | 434 | F | D | G | L | N | L | E | S | I | D | L | A | H | N | K | I |
|  | 458 | S | G | L | K | N | L | K | D | L | S | I | N | D | N | E | I |
|  | 482 | Q | S | L | E | K | L | K | K | L | S | L | L | N | N | K | I |
|  | 506 | K | D | L | K | R | L | N | K | L | K | L | S | S | S | G | I |
|  | 529 | F | D | G | L | T | I | K | S | L | D | L | S | K | N | G | I |
|  | 553 | S | G | L | N | G | L | K | I | L | S | L | R | D | N | R | L |
|  | 577 | Q | F | L | I | N | L | T | S | L | D | L | S | G | N | K | V |
|  | 601 | Q | R | L | A | R | L | H | N | L | N | L | K | S | V | G | I |
|  | 624 | F | N | D | L | S | V | D | S | I | Q | L | S | D | N | N | I |
|  | 648 | I | G | V | K | K | L | H | K | L | D | L | S | N | N | G | I |
|  | 672 | E | S | L | S | E | L | T | E | L | N | L | N | N | N | K | I |
|  | 696 | K | G | L | I | S | L | Q | D | L | S | L | V | S | A | G | I |
|  | 719 | L | D | L | P | N | L | T | K | L | D | L | T | K | N | N | I |
|  | 743 | Q | S | L | Q | N | L | N | Y | L | Y | L | S | E | N | K | I |
|  | 767 | K | K | L | S | S | L | N | S | L | Y | L | S | S | N | Q | I |
|  | 791 | A | G | L | D | K | V | K | Y | M | H | L | N | N | N | K | L |
|  | 815 | I | G | A | S | E | I | T | E | L | D | L | S | S | N | S | I |
|  | 839 | S | N | M | T | K | L | R | D | L | N | L | R | E | N | K | I |
|  | 863 | K | G | L | V | D | L | S | D | L | Y | L | K | G | N | N | I |

| **Protein** | **L0**  **position** | **-5** | **-4** | **-3** | **-2** | **-1** | **L** | **x** | **x** | **L** | **x** | **L** | **+6** | **+7** | **+8** | **+9** | **+10** |
| --- | --- | --- | --- | --- | --- | --- | --- | --- | --- | --- | --- | --- | --- | --- | --- | --- | --- |
| ER_PM1-  var2 | 165 | K | N | M | K | K | L | K | H | L | E | F | Q | G | R | T | N |
|  | 184 | S | L | P | D | S | L | E | D | F | Q | L | S | G | G | N | Y |
|  | 208 | K | D | L | I | H | L | K | N | L | K | I | T | E | A | G | V |
|  | 232 | S | Y | L | G | N | I | T | E | I | D | L | S | N | N | K | L |
|  | 255 | N | N | L | P | N | L | K | I | L | S | L | E | N | N | G | I |
|  | 279 | S | G | S | P | N | C | Q | I | I | H | L | E | F | N | K | I |
|  | 303 | A | N | L | R | N | L | S | A | I | D | L | S | N | N | K | I |
|  | 327 | E | G | C | S | N | L | N | S | L | N | L | D | N | N | L | I |
|  | 351 | K | D | L | I | N | L | N | Y | L | F | L | Q | Y | N | K | L |
|  | 375 | K | N | L | V | N | L | K | T | I | D | V | S | Y | N | E | I |
|  | 398 | F | D | G | L | N | L | E | S | I | D | L | D | H | N | K | I |
|  | 422 | S | G | L | K | I | L | E | N | L | S | L | N | D | N | E | I |
|  | 446 | Q | S | L | E | K | L | N | V | F | N | L | L | R | N | K | I |
|  | 470 | K | G | L | K | S | L | T | T | I | D | L | S | S | S | G | I |
|  | 493 | F | E | G | L | T | I | T | S | L | D | L | S | Q | N | R | I |
|  | 517 | S | G | L | K | T | L | E | D | L | S | L | N | D | N | E | I |
|  | 541 | Q | S | L | E | K | L | N | D | F | N | L | K | N | N | R | I |
|  | 565 | K | G | L | T | S | L | T | R | L | D | L | S | S | T | G | I |
|  | 588 | F | D | G | L | T | I | T | S | L | D | L | S | Q | N | R | I |
|  | 612 | S | G | L | K | T | L | E | D | L | S | L | N | D | N | E | I |
|  | 636 | Q | S | L | E | K | L | N | D | F | Y | L | T | N | N | R | I |
|  | 660 | K | G | L | T | S | L | T | R | L | D | L | S | S | T | G | I |
|  | 683 | F | D | G | L | T | I | T | S | L | D | L | K | K | N | N | I |
|  | 707 | Q | T | L | S | H | L | K | Y | L | Y | L | S | D | N | K | I |
|  | 731 | K | G | L | S | S | L | Y | R | L | Y | L | S | S | N | Q | I |
|  | 755 | A | G | L | D | N | V | K | Y | M | Y | L | N | K | N | K | L |
|  | 779 | I | G | A | S | N | I | T | S | L | D | L | S | S | N | N | I |
|  | 803 | S | D | M | S | K | L | S | N | L | N | L | K | E | N | K | I |
|  | 827 | K | G | L | V | K | L | R | Y | L | Y | L | K | G | N | N | I |

| **Protein** | **L0**  **position** | **-5** | **-4** | **-3** | **-2** | **-1** | **L** | **x** | **x** | **L** | **x** | **L** | **+6** | **+7** | **+8** | **+9** | **+10** |
| --- | --- | --- | --- | --- | --- | --- | --- | --- | --- | --- | --- | --- | --- | --- | --- | --- | --- |
| ER_PM1-var3 | 114 | F | S | G | V | K | M | K | E | F | H | C | S | S | C | R | F |
|  | 141 | E | D | Y | L | E | T | I | N | I | D | L | N | N | E | N | I |
|  | 163 | K | N | M | K | K | L | K | K | L | V | I | E | G | G | K | N |
|  | 182 | I | L | P | G | S | L | E | E | L | E | M | S | N | G | Q | Y |
|  | 206 | K | D | L | V | N | L | K | K | L | K | I | T | N | A | K | L |
|  | 230 | S | N | L | L | S | V | T | K | I | V | L | S | N | N | G | L |
|  | 253 | N | N | L | P | K | L | E | T | L | L | L | K | K | N | K | I |
|  | 277 | S | G | S | Q | N | V | I | Y | I | N | L | K | K | N | Q | I |
|  | 301 | S | N | L | K | N | L | L | K | L | D | L | S | K | N | K | I |
|  | 325 | E | G | C | S | K | L | I | K | L | K | L | E | K | N | V | L |
|  | 349 | K | D | L | T | H | L | Y | D | L | S | L | E | S | N | K | L |
|  | 373 | Q | N | L | V | K | L | R | K | F | K | A | K | D | N | E | I |
|  | 396 | F | H | G | L | N | P | Y | L | I | D | L | G | H | N | K | I |
|  | 420 | S | G | L | K | E | L | N | D | L | Y | I | N | D | N | E | I |
|  | 444 | Q | S | L | E | K | L | N | N | F | N | L | Y | H | N | K | L |
|  | 470 | S | N | T | N | T | V | T | S | L | Y | L | P | S | S | G | I |
|  | 493 | F | D | C | P | S | V | V | Q | L | H | L | N | K | N | K | I |
|  | 517 | S | G | L | N | K | L | S | S | L | E | L | S | D | N | G | I |
|  | 541 | Q | S | L | P | E | L | K | Y | L | H | M | S | G | N | K | M |
|  | 565 | T | G | L | K | N | I | Q | N | L | K | L | Q | S | I | G | I |
|  | 588 | F | D | G | L | T | L | S | D | L | D | L | S | E | N | K | I |
|  | 612 | S | G | L | S | G | L | L | S | L | N | I | E | N | S | G | V |
|  | 636 | Q | S | L | I | N | L | T | K | L | E | M | S | G | N | K | V |
|  | 659 | F | K | G | L | N | L | P | K | L | I | L | R | S | L | G | I |
|  | 682 | F | N | G | L | S | A | Q | S | I | D | L | S | S | N | N | I |
|  | 706 | S | G | V | K | H | L | N | S | L | V | L | K | N | N | N | I |
|  | 730 | E | S | L | S | E | L | H | K | L | N | L | N | S | N | K | I |
|  | 754 | N | G | L | N | S | L | Q | D | L | S | L | I | S | A | G | I |
|  | 777 | L | N | L | P | N | L | T | S | L | E | L | R | K | N | K | I |
|  | 801 | Q | S | L | P | K | L | K | S | L | S | L | D | G | N | K | I |
|  | 825 | K | A | L | S | N | L | D | K | L | D | M | S | S | N | Q | I |
|  | 849 | E | G | L | D | N | V | K | D | I | N | L | R | K | N | K | L |
|  | 873 | I | G | A | P | N | V | T | H | L | D | L | T | H | N | S | I |
|  | 897 | S | D | R | P | K | M | Y | S | L | N | L | K | D | N | K | I |
|  | 921 | K | G | L | T | S | L | R | H | L | T | L | E | D | N | N | I |

| **Protein** | **L0**  **position** | **-5** | **-4** | **-3** | **-2** | **-1** | **L** | **x** | **x** | **L** | **x** | **L** | **+6** | **+7** | **+8** | **+9** | **+10** |
| --- | --- | --- | --- | --- | --- | --- | --- | --- | --- | --- | --- | --- | --- | --- | --- | --- | --- |
| ER_PM2 | 101 | F | S | D | A | K | I | K | R | F | H | C | V | D | C | D | L |
|  | 126 | P | L | E | N | I | L | E | E | L | K | L | D | D | C | T | I |
|  | 148 | Q | N | L | K | T | L | K | S | L | K | I | D | S | D | N | L |
|  | 171 | V | I | P | S | P | L | E | H | F | A | I | Y | N | A | K | F |
|  | 195 | T | T | Y | S | S | L | T | T | L | K | I | Q | S | T | N | L |
|  | 219 | Q | S | L | K | N | L | T | L | I | D | L | S | M | N | R | L |
|  | 242 | K | N | L | P | E | L | Q | E | L | F | L | E | Q | N | R | I |
|  | 266 | T | N | M | K | K | L | K | F | I | D | L | D | M | N | E | I |
|  | 290 | S | N | L | P | N | I | V | S | I | A | L | G | E | N | N | I |
|  | 314 | L | G | A | D | E | I | K | E | L | N | L | G | N | N | E | I |
|  | 338 | A | K | L | A | K | L | E | R | L | N | L | A | V | N | K | I |
|  | 362 | S | D | C | L | N | L | M | D | L | N | L | K | S | N | N | L |
|  | 386 | S | H | L | K | K | L | K | R | I | D | L | W | H | N | G | I |
|  | 410 | D | G | L | L | K | L | T | S | F | S | L | V | S | N | S | L |
|  | 434 | E | G | L | E | S | V | E | I | L | D | L | H | D | N | K | M |
|  | 458 | S | D | M | P | K | L | H | T | L | L | I | N | N | N | K | I |
|  | 482 | K | D | L | P | N | I | D | H | L | N | L | N | F | N | A | M |
|  | 506 | V | G | A | S | M | L | K | E | L | D | L | E | E | N | P | I |
|  | 530 | H | G | L | E | S | I | V | K | I | S | L | S | K | A | K | L |
|  | 554 | S | G | L | P | S | L | T | T | F | R | C | F | S | S | D | L |
|  | 578 | K | S | T | N | E | L | H | D | L | Y | L | R | Y | N | K | I |
|  | 602 | K | G | L | S | S | L | E | D | L | D | L | A | Y | N | N | I |
|  | 626 | M | G | L | S | V | L | H | K | L | D | L | S | W | N | K | I |
|  | 650 | L | G | L | T | N | L | K | I | L | Q | L | D | N | N | E | I |
|  | 674 | N | G | L | E | S | L | E | T | L | D | L | S | H | N | R | I |
|  | 697 | F | E | F | S | N | L | Q | V | L | E | L | E | S | N | L | L |
|  | 721 | D | K | A | P | T | L | T | S | L | N | L | H | G | N | G | I |
|  | 745 | K | N | L | V | N | L | D | K | Y | H | V | L | T | S | K | T |

| **Protein** | **L0**  **position** | **-5** | **-4** | **-3** | **-2** | **-1** | **L** | **x** | **x** | **L** | **x** | **L** | **+6** | **+7** | **+8** | **+9** | **+10** |
| --- | --- | --- | --- | --- | --- | --- | --- | --- | --- | --- | --- | --- | --- | --- | --- | --- | --- |
| ES_PM1 | 63 | L | K | N | L | T | N | E | E | F | D | F | S | G | T | K | M |
|  | 98 | C | N | F | T | G | V | T | S | A | L | F | T | S | S | E | S |
|  | 108 | F | T | S | S | E | S | T | L | L | S | V | S | S | Y | S | S |
|  | 134 | K | N | L | K | K | V | K | T | L | S | I | R | S | Y | N | K |
|  | 159 | D | L | P | P | S | L | E | S | I | D | V | N | S | L | M | A |
|  | 183 | N | F | L | P | N | V | T | N | L | A | I | T | Y | S | L | I |
|  | 207 | E | N | F | P | N | L | K | T | L | Y | L | N | Y | N | K | I |
|  | 230 | K | E | M | S | K | L | K | S | L | D | I | S | Y | N | N | I |
|  | 254 | S | G | L | E | N | L | R | R | L | I | L | N | S | N | P | I |
|  | 278 | Q | K | L | S | N | L | K | K | L | E | I | R | G | S | K | I |
|  | 302 | K | G | L | K | S | L | T | S | I | S | F | S | T | P | S | L |
|  | 326 | E | D | L | N | N | V | S | S | L | S | I | K | Y | G | H | L |
|  | 350 | N | N | L | K | N | L | E | S | L | S | I | K | N | Q | Q | I |
|  | 373 | F | N | G | L | S | I | D | F | L | D | L | S | D | N | K | L |
|  | 396 | S | H | L | E | S | V | K | K | V | D | L | S | R | N | L | I |
|  | 420 | D | N | V | P | H | L | D | T | L | F | L | S | N | N | K | L |
|  | 444 | Q | N | L | P | A | L | S | Y | L | D | L | S | S | N | E | I |
|  | 467 | F | S | G | T | N | I | K | T | L | L | L | D | E | N | R | L |
|  | 489 | S | G | L | D | N | L | E | V | L | N | I | R | K | N | N | I |
|  | 513 | N | S | L | H | K | L | K | D | L | T | L | Y | G | N | K | L |
|  | 537 | K | G | L | N | D | L | T | H | I | N | L | R | N | S | S | I |
|  | 561 | N | G | L | N | K | I | E | S | L | S | L | A | Q | N | Q | L |
|  | 585 | L | G | L | K | Y | V | N | N | L | D | L | S | S | N | Q | L |
|  | 608 | F | D | S | L | P | L | S | E | I | N | L | Q | N | N | Y | L |
|  | 630 | K | D | L | K | S | L | K | V | L | D | L | S | S | N | K | L |
|  | 653 | L | N | Q | I | L | L | E | E | L | N | L | Q | N | N | N | I |
|  | 675 | E | L | S | G | N | L | K | K | L | C | L | S | G | N | P | L |
|  | 700 | K | M | G | T | N | R | K | E | L | I | F | N | R | N | C | F |

| **Protein** | **L0**  **position** | **-5** | **-4** | **-3** | **-2** | **-1** | **L** | **x** | **x** | **L** | **x** | **L** | **+6** | **+7** | **+8** | **+9** | **+10** |
| --- | --- | --- | --- | --- | --- | --- | --- | --- | --- | --- | --- | --- | --- | --- | --- | --- | --- |
| EB_PM1-  var1 | 109 | D | S | N | S | S | L | N | S | F | E | I | T | A | D | N | M |
|  | 132 | K | N | L | K | N | I | T | S | I | Q | L | T | L | S | T | D |
|  | 157 | D | F | P | S | S | L | M | K | L | E | L | L | Y | F | K | L |
|  | 181 | N | P | L | P | N | L | E | Y | L | Q | L | N | N | D | E | I |
|  | 205 | N | N | F | P | N | L | K | E | L | D | L | Q | S | N | K | L |
|  | 227 | S | G | L | P | H | L | V | K | V | D | L | S | S | N | D | I |
|  | 251 | T | D | L | K | N | L | K | N | L | N | I | E | H | N | K | F |
|  | 275 | V | E | L | K | S | L | T | N | L | K | I | G | N | R | F | L |
|  | 299 | D | G | L | N | N | L | E | S | F | D | L | T | S | T | G | Q |
|  | 324 | N | N | L | K | K | I | T | L | L | K | L | N | N | M | D | L |
|  | 347 | F | D | G | L | N | V | A | K | I | E | I | M | W | S | K | I |
|  | 371 | S | N | L | L | Q | L | K | S | L | I | L | S | H | N | K | I |
|  | 395 | G | N | L | P | K | L | T | N | L | Y | L | G | F | N | E | M |
|  | 419 | Q | S | L | S | A | L | N | R | L | E | I | D | S | N | N | I |
|  | 442 | F | V | G | T | N | L | T | I | L | D | L | S | Y | N | K | L |
|  | 465 | S | G | L | E | H | L | M | T | L | D | L | S | N | N | K | I |
|  | 489 | H | N | Y | P | N | L | E | I | L | K | L | S | G | N | K | I |
|  | 513 | Q | E | L | I | K | L | K | Q | L | E | L | M | N | S | N | I |
|  | 537 | N | G | L | K | N | L | E | D | L | D | L | S | Q | N | L | I |
|  | 561 | E | A | T | P | N | L | K | K | L | N | L | D | Q | N | Q | L |
|  | 583 | S | N | L | K | N | L | H | S | L | K | L | S | S | N | K | L |
|  | 606 | V | A | M | P | S | L | K | T | L | D | L | T | S | N | D | I |
|  | 628 | E | L | N | R | N | L | T | D | L | C | I | S | G | N | A | I |
|  | 649 | S | F | Y | K | N | I | L | S | G | Y | C | T | D | F | P | I |

| **Protein** | **L0**  **position** | **-5** | **-4** | **-3** | **-2** | **-1** | **L** | **x** | **x** | **L** | **x** | **L** | **+6** | **+7** | **+8** | **+9** | **+10** |
| --- | --- | --- | --- | --- | --- | --- | --- | --- | --- | --- | --- | --- | --- | --- | --- | --- | --- |
| EB_PM1-  var2 | 76 | E | L | V | K | N | L | S | R | P | E | I | S | F | W | G | N |
|  | 100 | F | A | G | I | K | V | R | K | F | H | C | F | D | C | N | Y |
|  | 125 | D | S | V | T | S | L | T | S | V | D | D | D | S | R | D | M |
|  | 148 | E | K | L | K | N | L | T | S | L | K | L | T | F | S | T | D |
|  | 176 | I | L | P | Q | T | L | K | K | V | T | L | K | H | F | K | L |
|  | 200 | N | S | L | T | N | L | E | Y | L | N | L | Y | S | D | D | I |
|  | 224 | S | N | F | N | N | L | K | E | L | K | L | G | K | N | K | L |
|  | 246 | S | G | L | H | Q | L | E | K | L | D | L | S | N | N | H | L |
|  | 270 | T | N | L | E | N | L | K | E | L | N | I | N | S | N | R | F |
|  | 294 | S | G | L | K | S | L | I | R | L | D | I | K | Y | T | S | L |
|  | 318 | D | G | L | K | N | L | A | S | L | E | I | I | D | T | G | T |
|  | 343 | H | D | L | K | K | M | I | S | L | K | L | I | N | M | Q | L |
|  | 366 | F | D | G | L | N | V | A | E | I | E | I | S | N | N | K | I |
|  | 390 | L | N | L | Q | R | L | T | S | L | E | L | T | N | N | Q | I |
|  | 414 | N | G | L | H | S | L | T | Y | L | N | L | A | N | N | H | I |
|  | 438 | Q | S | L | S | A | L | E | T | L | D | L | S | G | N | S | I |
|  | 461 | F | I | G | T | N | L | T | N | L | I | L | I | H | N | K | L |
|  | 484 | S | G | F | E | H | L | L | S | L | D | L | S | D | N | D | I |
|  | 508 | H | N | Y | V | N | L | E | S | L | N | L | N | D | N | K | I |
|  | 532 | Q | D | L | I | N | L | N | K | L | T | L | D | D | N | N | M |
|  | 556 | K | G | L | K | N | L | K | D | L | G | I | P | F | E | L | R |
|  | 580 | E | M | G | P | N | F | E | Q | V | N | F | V | D | N | F | D |

| **Protein** | **L0**  **position** | **-5** | **-4** | **-3** | **-2** | **-1** | **L** | **x** | **x** | **L** | **x** | **L** | **+6** | **+7** | **+8** | **+9** | **+10** |
| --- | --- | --- | --- | --- | --- | --- | --- | --- | --- | --- | --- | --- | --- | --- | --- | --- | --- |
| EB_PM1-var3 | 74 | F | T | G | I | E | L | Q | E | F | E | C | T | D | C | N | Y |
|  | 99 | D | S | N | S | S | L | T | S | F | K | V | T | A | A | N | M |
|  | 122 | K | K | L | Q | N | I | T | S | I | R | L | T | L | T | T | D |
|  | 147 | N | F | P | S | T | L | K | K | L | K | L | K | E | F | K | L |
|  | 171 | N | P | L | P | N | L | E | Y | L | Q | L | Y | N | D | E | I |
|  | 195 | N | Y | F | H | N | L | K | E | L | D | L | G | S | N | K | L |
|  | 217 | S | E | L | S | Q | L | E | N | L | D | L | S | Y | N | E | I |
|  | 241 | S | N | L | K | N | L | K | N | L | N | I | Y | H | N | K | F |
|  | 265 | A | E | L | K | S | L | T | K | L | K | I | S | N | R | F | L |
|  | 289 | Y | G | L | D | N | L | E | S | F | A | L | S | F | T | G | Q |
|  | 314 | N | N | L | K | K | I | T | S | L | R | L | N | N | M | E | L |
|  | 337 | F | D | G | L | N | V | E | E | I | K | V | T | N | N | K | I |
|  | 361 | S | N | L | P | Q | L | K | S | L | R | L | S | S | N | E | I |
|  | 385 | G | N | L | L | K | L | T | Y | L | N | L | E | F | N | K | M |
|  | 409 | Y | M | L | S | T | L | N | E | L | D | I | S | F | N | N | I |
|  | 432 | F | V | G | T | N | L | T | I | L | D | L | S | D | N | T | L |
|  | 455 | S | G | L | K | H | L | V | T | L | D | L | S | N | N | E | I |
|  | 479 | H | N | Y | P | N | L | E | I | L | K | L | S | G | N | K | I |
|  | 503 | Q | H | L | I | S | L | K | Q | L | E | L | I | N | N | K | I |
|  | 527 | N | G | L | K | N | L | E | D | L | D | L | A | N | N | L | I |
|  | 551 | E | P | T | H | N | L | K | K | L | N | L | D | N | N | Q | L |
|  | 573 | S | N | L | K | N | L | H | S | L | K | L | S | S | N | K | L |
|  | 596 | I | A | M | P | S | L | K | T | L | D | L | S | S | N | N | I |
|  | 618 | E | L | N | R | N | L | T | N | L | C | L | N | G | N | A | I |
|  | 639 | N | F | Y | R | N | I | I | S | F | D | C | D | D | S | P | V |

Table S4. Sequence homology search of LRR proteins against NCBI non-redundant protein using BLAST.

|  | **Protein name** | **Accession** | **Species Name** | **Query Cover** | **E-value** | **Per. Ident** |
| --- | --- | --- | --- | --- | --- | --- |
| ER_PM1-var1 | insulin-like growth factor-binding protein complex acid labile subunit | XP_064644154.1 | Lineus longissimus | 85% | 1E-94 | 31.9% |
| ER_PM1-var2 | insulin-like growth factor-binding protein complex acid labile subunit | XP_064644154.1 | Lineus longissimus | 79% | 4E-95 | 32.4% |
| ER_PM1-var3 | chaoptin-like | XP_044005686.1 | Aphidius gifuensis | 88% | 1E-97 | 31.2% |
| ER_PM2 | insulin-like growth factor-binding protein complex acid labile subunit | XP_044019265.1 | Aphidius gifuensis | 87% | 1E-80 | 29.1% |
| ES_PM1 | chaoptin-like | XP_044005686.1 | Aphidius gifuensis | 78% | 9E-66 | 36.3% |
| EB_PM1-var1 | leucine-rich repeat-containing protein 15-like isoform X8 | XP_056005023.1 | Ostrea edulis | 67% | 9E-51 | 35.1% |
| EB_PM1-var2 | leucine-rich repeat-containing protein 15-like isoform X13 | XP_056005028.1 | Ostrea edulis | 73% | 1E-48 | 34.1% |
| EB_PM1-var3 | chaoptin-like | XP_044005686.1 | Aphidius gifuensis | 73% | 4E-48 | 31.6% |

Table S5. High-MW protein sequences

**>ER_PH1 (accession: PP551269.1)**

MKTIILVVAFLLVVEGATRHKSKTTHKGGQRGGYGGVGGHGGGYGRGGKGGRVRGSGGGGGDYGGGGGNYDGGLDEGFGGGGYGGSSGGGSRGGSGGGSRGGSGGGSRGGSRGRSRGGSGGGSGGSSGGGSGGSNGGALSYEEYEKLKQNPQITKGSNPVLSLGQPNHRSNFKINQNPNNAYDFDINPDGNDPENPEIIPQGDKGKPDYSVILGVGPKNNRKYVRMAPKAGQPGEFDFDPVTMGPSTPAEEDSNPPDNGGSEPGQNGATRIPQDEYERLKQNPQITQKGNNKILSLGQPNHRRHFKINQNPNNPNDFDVKPIGSSDDPNSEPNENDPEKPEIIPQGNNGKPDHSVILGVGPKNNRKHVQMKPKSGHPGEFDFDPVLMGPSTPVDDEDHPDDGGPDLVVKVPEDEYERLKQNPQVKSKGKYRILSLGQPNARRHFKIKPNRKNPFDFDVEEIGDPDDPDSKPDSNEAEKPEINPENRKGEPNYDVVFGVGPKNDRKYVRMTPDPKKFGKYNFEPVLLKPGDSFHPQTPEPTPAPEPEPETTTAPTPTTTEQPEEDEPEFEDCLEDPTFRNVRAPGGKRPYQIISVGPKDHRQHLKKTPRSNNPNDYDLEPYDPNTPNHRPNPKDSYAPEVFPNGKRGTPDYSPVINTGPKDHRRPYRLVPHPENPNHSGFIPVKRGPKKPGKKHPRFVDIPRKTRPRKKGEPKPEEDIPEDGEPEFDDLPEDPEFRTRKTPGAKKPYTLISVGNPPNRRTFKKTPGNPDNPDDYILEPYDEDTPDHKPDPDDPSAPKVYPNGKKGKPDYSPVIEIGPVNNRKTFRLTPHPTNPNRVGFIPVRKLPGKKHPRFRDIPRKLRPHGYRPASTPEPNDGDDVFEDLPDDPEVRPHNRPGEKYPSQIISVGKPPYRRHFEKKPKPEDPKKFDLVPYNPRKPDEKPDPENPNTPKVYPNPKGGDPVIRTGTPEHPKYHKVVPDPKHPDDPEKVNFVPVEPVGDPDDDGEPTFEDKPRKSHKYHTTAKPKKTTAHPNDGDPIFEDVPEDYDIRTIKKPGSKHPVKLISIGKPPHRQQFLKEPGPSGRPDDFTLTPYDSNKPDNKPNPEDPYTPEVYPGKPPVIRVGPKDDPEYFELKPDPKDPKNPNKVTFIPVKPKPHKPGQPTRFVPKLPRRFHPYGKRTTTQPQDGDIEYTDGPDDPEVTTLPRKPGQKYPSQLIAVGKPPNRKYFEKRPGKSGRPDDYKLLPIDPNNPNKPLDPKNPKTPKVYPGIGKPGTPNYEAPVIETGTPKNPKFFRVVPNPNEPNNPENVEFVPVQKVKKPHRGSKFQPKPRHTHRYRTTKHPRPRPDQTPAPIDGEPVFVDEPDNPKVYRVKVPGEKRPREIIGVGKPPNRKFFEKIPNRKNPKQFTLRPFDPKHPGKTPDNCKVYPPKPGKDGPHNPVIEIPTDDGPKFYEVKPGPKFIPVEKLPGDEFKRLPRITRIYRRTTKKSKRHTTTPEPNENEIVPDDGPEDPKFYPFHEPGSKRPSKLIAVGKPPYRRIFKLVPGPGDKYTLKPFTPKGKKTPKDTQVYPNNGTPVFRVPDKKGHPKYYKLVPDKKHPGKFDFVPVKPLSPKGNKFNPLPRQTRHYHTTKKHRRTTTPEPDEGTPIFVDEPFVEYRPYKRRPGQNYPSQIVTVGKKPKVKYFEKIPSNENPKEFTLVPFDPHRPNKPPKKGVIPPTDDRKYPVIEGPDKKFYVCKPNPKNPNEITFVPVVPDGDNFEEVPRVWYPYHTTRNPEEVELGISTTPSEDNTEPSSGNTTPDDIGGLETATPEDTEAVLLETTPTPTEIEATTDELSEGTTEPEITPEILDTEKPEEPSDVNDSREPTAPGTDYGSGDNTPSGTPEIEGPEINKPGSPPGSTNVYKPDHPEIPGEPEETPEAPGPNVEKGETPEGPEPNGGPGETPDIEKPHGEPGETPGKPEPGETPAPGETPAPGEKHAPGETPAPGETPAPEETPAPGEKPKSPGDINKYCMKKYENCKTRGY*

**>ER_PH2 (accession:** **PP551270.1)**

GDGGESLDGYVLQPDGSYLKTITDDQPEMNYQQDGNLPPGFVAPPGFSGQDGGWILQPDGTYMKTIVETEAPVEETVEIEPDYQPDEFFQVGQPGAPNSYMIFGFGEPNQRNYFKQTPGGSGDPEDYSLEPVVSPSNPVPVNSKDRNSGRVIRKGKRGEPNYKTVVQVGGPRNRRFYRLVPDSSKKSGYKFIPQRATGSLKKPGGSTQPNKVPEEDEPEQFSEETEEVEEQEEEDDDMPDDPTIDQMDGYQLICIGKPNRRYLKLTRPRSNNPYDFSLEPIKSPDSPNDKPDPNDDEAVEVIPTGEQGKSSYRPIIAVGPKYNRRYVSVNQDPEDPDSFTFRPVKRRLRGPGETQTEPNDDDDDVSYDDIPENPEYKTVAAPGQKTTTEILALGPKEHRIYFKIIRKKPSDPNDVDIIPIQDPEHVDSPPNKDDPYSPKVVANGEPGESGDGPIIAVGPKGKRVYYRLSFKKKGLPKFIPLRKKKRPGKLRPIFSKKPRQWHWLTKHKLSNKPAQPSHPVFEPQKPIFSNKPKVSQVKTVKKPGIKEPFEVITLGPATKKVYIKKTPGPSGKPTDYKLEPLGNPDDPNSKPNPNDPDAPEVIENGKPHTPDYSPVVAHGPKNNRKYVIVKPHPADPKRVKIHPAEILPGSDPKNPKFKISVKKTTPIFRPKPHKPPIGIHPNFTPHLPNWLDLLTGIRRPIPGLPPFSHVRFPPVKFIRKPEQVSITITLPPIEVKVNHKPGDKNPYQVISFGPKDKQIHIKKTPGPSGHPNDYKLEPLGNPDDPNSKPDPKDPDAPEIIENGKPGTPDYSPVIAHGPKNNRQYTHLQPHPKDPTKINIKTAELVPGSDPKKPTFKPLNSKGPLIRPRKPWINPGFLRPMIDPGFSRPMIIPGFPRQIDPGFAGPSIDPGFLRPLRPERIPVEVKVNHKPGDKNPHQVISFGPKGKRIHIKKTPGPSGDPNDYKLEPLGNPDDPNSKPDPKDPDAPEVIENGKPGTPDYSPVVAHGPKNNRKYAIVKPHPSDPKDLSRVKLRPAEIVPGSNPKKPTFRVKPRRTPERPVFGDRYKPEVKDSKPGDKNPHQVISFGPKDKRIHIKKTPGPSGHPNDYKLEPLGNPDDPNSKPDPKDPDAPEIIENGKPGTPDYAPVVCHGPKNNRKYVILKPHPSNPKDVKLTPAELAPGSDLKKPIFRIKPRRLTLPTKPSYNPFDDDSEQPEVKDNNKPGDKNPYQVISFGPKDKKIHIKKTPGPSGNRNDYKLEPLGNPNDPNSKPDPKDPDAPEVIENGKPGTFYYAPVVAHGPKTNRKYTMLLPHRKDPKLIKLEPAQIDIGSDPKKPTFSRSFKYGRGFRKHKFNPVCYRQYSECNERSE*

>**ES_PH1 (accession: PP083421.1)**

MKILLSVLVLLIVVECGNSRKIRHRGGSRRGSGGGSGGSSGGSSGGSDGSYGGSDGGSGGSYGDSGGGSGDSTGSNGGPGDSYSESGGSSGDGGSGGSYGGSDGGPGGSYGGSGGSGGGGGGGGGGGGGGSGSDNNPPEGYYDPNKSGPLPPGFEAPPGYEGGEWHPGPDGTMVRTIVEEEPGTETTELVPPDYDPQITPVGQPGSPGYHDIIGVGKPGERGYFKRTPGPSGNPDDYTLEPIKSPENPEPVDPNDPNAPRVIRTKKNKNPTLQVGNKDNPQYFGLKPDPKNPGHFTLVPRKMHPGKPGHRKPGHPKGGKGKPGRGKPGHGKPGDGGALSPEEYEKLKQNPVMKYPVFTLGKPEHRRSFRITQDPNNPNDFSVDPIGNENDPQSPADPNDPEAPEIIPQGEKGKPDYNVVIGVNPKKKRDYVQMKPDPKNPQQFNFDPIFMEPQQEEPQQEQPTPAGGEPEVFLDPGSESVTPEEYERLKQNPKIDRKKGSPILHLGYEPYRRPFRIKQNRRNPLDFDVQPIGSDNDPNSKPDKDDPYKPVVYPLGEKGQPDYSQIIGVGPKDKRNYVRMIPDPKKPGKFEFEPVLLKPGEPIPFVTETEPPPTPTPEPTVPPPPPTHPPTTKKPEEEEPCFEDLPEDPEFRRVRHPGRRRPYQIISLGPKDHRQHFKKTPKSNNPDDYDIEPYDPDTPDHKPKADDPYAPEVIPQGQPGTPSFEPILMTGPKDNRKPYKLVHHPENPSRSSFVPVRPLPKKPGQKHPKFVNAPRQIRPRKKDEPRQPDDEPEEGEPEFTDLPDRPRFKHVKQPKRPYTLISVGKHPHRRTFKKTPNPNGNPDDYDLEPYDEDTPDHKPNPHDPSVPKVHKNGKKGTPDYQPVIEIGPINKRKLYKIKPHPSNPNKVEFVPVKKIMRKGKPFFKELPRRTATKRFRPRTTPKPNEDDDVPDDLPYDPEVRPHRTPGQKYPTQIISVGKPPNRKHFKKIPKPEDPKKFDLEPIDPDDPDEPLDPEDPNTPKVYPNPEGNPVIETGTPDDPKHHEVNPDPEDPEDPERVSFTPVKRLGEPGEKNPKFEKIPRKSHPFHTKRPEEATAPEPEENEPVFEDIPYNPTFKTLHRKRPDRPAQIIALGRPPHRQEFIKEPGRSGKPHDFKLSPYDSRRPDKKPDPNNPNKPEVYPARGKPGKPGYQPPVIKTGPKDDPEYFEIHPDPSDPKNPNKVLFVPVKARPGKPGTRPKFSPKEPRRYHPYHDPLKPRPTTEEPEDGTEYFVDQPEDPEVTTLFRPLRKYPSQLIAVGKPPNRKYFEKKPGPSGHPKDYTLHPIDPKHPDKSLNPKDSKTPKCFPGEGKPGTPNYKPPVIQTGSPDNPKFYRVIPNPKDPNNPESVEFIPVKKDKKNPKKFKPLPRNVRIYHTTKKQATSQPNQTPPHEEGDPYFTDLPDNPVVLTVRKPGSKKPVQILALGKPPHRKYFEKVPEPGKPDRFTLKPINPNGKPGEPSKVYPGGNKPGSRYPVIEYGKPPKAFEVRPDPLHPNDPTKVQFVPVKKVPGTDGPSFVPAPRTVGKYHTTRKKHHKTTKEPKEGSVIYDDGPYLPKFYPGKKNTKIVAVGKPPKRRYFKIIPKPDKKFTLKPIDPQNPDDPNPHPEDTKVYPQPDGHPVFEIPGPNKKPEYFKLVPKKDNPKEMDFIPVRPVSPGSFTSKPRTTRHYKTTKKSKRTTRPTSRVIFEDIPHNTQVPKKRPNQEHPSTIITVGNEPNQKHFEVIPDPSGNPKKFTVVPYNPYHPNKPCDPKNPKTPQVFEVPKDYPVIQGPDKKFYRLEPDKHNPGQYTFVPVTPLDENHKPSNPHKPDVSFVPIPRKWHLRTTPGIDDWLDLSTTPSPKTTTDNLGTGYSVVTKPEEISEPTTEVDNAETTVGEEELNKEPTEGPMTTEATEQFSTEQPTVPSHTETEVAIELTTAEEIILSTASVTQPTVAGKTESTPAELTGKSEIPQSTVSTTEAPTVATHTPMSVTTEAIQGTTQGAGVSTESETKGQPTGATSGPEEPSKQEPTTSEVSGTGQTASPATSSEAHPRPTAPNVDDVCFKAYDDCKRSRSSYA*

>**ES_PH2 (accession: PP083422.1)**

MEMMYTLFFLLFGIVHGQGDGWVLQPDGSYMSYGDGSSGGSYGSTGGSYDGSGGLYGGSSGGSYGELGSGGLFGGSGGGGFGPGGSYGGFDGGLGGSSGGSQGLPGNGWILQPDGSYLKYEGSGGGGGGGGGGGGSGSDGPPGNGWILQPDGSYMKYDTSGEGNQGSAGSYGGNLPPGFVPPPGYSGNEGGWVLQPDGTYMKTIEQTEPPTEIRYPVNPDYQPDETLESGKKGSPGYSQVVGFGEPGNMNYFKITPGDAENLYDYSVEPVVSRDNPTHSTDTEHGKVISRGTKGEPDYQPILQVGPRNHRRYYKMLPDSSKPSGFNFIPQRLTGWLKKQGGDDTSGEVLEPEESEVTEEANEEIPDEPEIEKKNNDFIICIGRPNRRYLRLKRYSNNPHDYSVEPIGSHDNPDSKPNPNDEEAPEVMTEGTKGNPNYRSIIAVGPKYNRRYVEVLPNSNDPDKYSFAPVDVQVQPGEQATEPNEEDVEYDDLPENPEYKTIPGKKKSYEIFSFGPKNRRVYFRIIRRNPNNPNDIEVFPFKDPEHPDSPPSSDDPSAPEVIQQGDPNDPEDGPVIAVGPKNKKVYFRLGIRKDKVPKFTPVRRGKRIPGKHRLVFTKKTRIWHSRPQKKPLVRKPSAPLKFETSAPVEAATDYSPDEVDVNPENPVEKRPYQIITFGPPKKRIYIKKTPGPSGNPYDYKLEPIGNPEDPNSKPDPKDPDSPEVIETGEPGTPSFAPVVAHGPKQNRKYVIIRPNSENPKDPNQMEFSPAEPVPGSDPKKPRFRILRAAVPVAPKDQFTDNTPKNPNIKTALKPGDKTRYEVISFGPKNRRIYLKKTPGPSNKPDDFKLEPIGNPNDLNSKPDPNDPEAPEVIENGQPGTPDYSPVVAFGPKTRRKYIIVKPSPKNPKEIQFIPAEPEPGSDPKQRRFKPLTKKVLPGLIRPKPPTHPDSPEIKTVEGPHPHQVISFGPENKRIHVKKSPGPSGDPNDYKLEPLENPFDPNSKPDPQDKDAPAVIETGKPGTPNFLPIVSYGPHDNRKYMSFKPIGNTIALSPATLVPGSDPKHPAFNIKTTHPVSPQIPSILKDLHIPTHPLIPPIFNQLPPILNPRGINLLPGLLPVISNLAELTKPTVQNVEKPNKPSYQVLSFGPLLNRIHIKKTPGPSGKPDDFKLEPIKDPNDPDSKPDPKDPQTPEIIESGKPGTPDHQQVIAFGPKPQRKYVLVKLNPTDSKKLELSPVEPEPGSDNKQPKFRPVVKPVIPTLPLPSLDLHNPVINTVEKPNKPHYQVISFGPSTQRIHIKKTPGPSGKPEDYKLEPLGNPDDPDSKPDPKDPNAPEIVDSGTPGSPDYEPVVAYGPKTARKYVILKPVPLNPKALDLQPAVLEPGSDPKKKKFKVVIQPGISPLLPQILPPFWPKPSIPYLPKPPPITLPLISHGLQNPIVNNIEEPNKPHMQIISFGPILTRVHIKKILGPTGNPNDYKLEPIGNPDDPDSKPDSKDLDAPEIIENGKPGTPDYKPVVAYGPKSLRKYVTLQSRPSNPKEIDLEPAELAPGSDAKKPIFKYLPAIPVFGNKDRNAPEVKTVTDPISKKPVQVISVGPIFKRVHLKKIPGPSGKPNDYKLEPLGNPDDVNSKPDPKDPDSPEIIENGTPGTPDYAPVVAYGKKIRRKYLKIKSNPLQPNDPTKLLFNRVRLLSPKPGEKTPTFSESLIPKFNPVCYKQYQDCNDRS*

**>EB_PH1 (accession: PP551271.1)**

MKIVILLLAVLLVVQCEAKVRRRGSMRHGGGGSYGGGGGDGYGGSGGSYGGGSSSGGYGGDNGGSGDEYGGGSGGGDSGGGGGGEGGSDDGGSGGGGGGRGGGGGGHGGGGSGGSGGGGGGGGYGEGPEGGSGNNGGDSEALSPEEYERLKQNPHLNYPVMTFGTDHITPFRVTQNSDNPYDFNLDPIGNEHDPQSSADPNDPNTGEVYPQGNKGQPDYSIIVGVNPKGNNRKHVRVSPDPKKLHEFTYDPVNMESQGPEPQPDQDDFGPDTPEGGKRPGSGPEEKTTPSPKASIVGTGKTVSPEEYERLKQDPKIDKKKHPNILHLGYGPERRPFRVDQDPNNPSDFSLKPIGNENDPKSKPDKNDPYKPVAYPIGQKGQPDFSVVCGVGPKDNRNYVRVIPDPNKPGKFEFEPVLLQPGDPIPFPTDEPEATPEPTTTPPPTTTTKRTTTPQPEEEDECFDDLPENLQFRTIRRSGQKHPYQIIAMGPKEHPQRFRKIPRSNNPYDYDLEPYDPNTRNHKPSPDDPWAPEVIPQGKRGTPSFQPIIRTGPKDDPKYFRAIPHPQNPNRLAFYPVRPLPKKPGQKHPQFVNIPRKTRPRRKNEPKESDDDTPEEGDPDYDDLPKNPKFNRVKKGKKPYDIIELEDEPYRRKFKKTPNPNGNPDDYKVEPYDEDTPNHKPDPSDPGQPKVYPVGRRGKPDYSPVIEVGPPNRRKCFRLVPHPTNPHKVSFLPVKKIPRRGKPPLFQDLPRRVKPYRFRPKTTPEPNEDDEVPDDLPYRPEIRPYRRNPREKQLSVGRPPNRKHFRKIPKPDDPQKFDLVPTDPENPDEELDPDDPKTPKVYPNEDGGAPVIETLGPDKKPQYHEVHPDPDDPNDPEKVSFIPVKPVGDPNQKNPKFETLPRRTRKFHTRRPKTTPKPQPGDPVYEDQPKNIVIKRIPRKKPKRPAQSIKVGPDNHRQEILKEPGPSGKPRDFKLSPRDPNKPDQKPDPSNKWTPEVFPPRGRPGQPGYQPTVVRFGPRDDPEYYEVHPDPSDPKNPNKVTFVPVRRLPGKPGTRPRFSKIPRRYHPYVDLRKPKPTTEEPEEGHPVYKDVPHYRVSTLFRPLNKYPSQKIEVGDEPNKKTFEKVPGKSGHPKDYTLRPLDPKHPNKPPDPKDPNTPKVFPGTGKPGTPNYKPPVIQTGSPKNPKYYRVVPDPEEPNNPEKIQLIPQKVTKRHGKKTFIPKVRIVHPYHTTKKAATHQPNQKPHEDGKPYYVDEPENPKVYNVEDPHSKKLVQLIAVGKKPHRKIFKKVPKPGNRYDLEPLTPKDKTDEPGKVYPGDDTHYPVIEYGKPPKTYEVRPDPEHPHDPTKVVFVPVTKVPGPTGPSFTPAKRKTKIYHRTTRRHSTTKEPKEGDIIYTDEPENLKSYPDHRPHQPKGPVVAMGRYPKRKYFKCVPKPDGKFDVKQFDPQDPDNPDPHHDDTKIYPQDDGHPVIQIPGRQGKPEYYKVVPDKHDPQKPSFVPVRPLSPGTFTKQPRHTKHYRSKKPKTKKTTTPEPEFYFRDVPPHNIRRRKRPGHPDSYILTVGKKPNEKTFELEPTDDPHTFNVIPFNIYHPNRPSNPKKPGTPRVYNVPGDNPVIQGPDKKIYRIVPDPKTPEKFNLVPLKCIKPGKPGEPEEPGDPNEPGVRFVPKPREWVPRTTTPDPDDAGLINKLTTPEKEEPTDAMPDGYTLLAQEPIETTTEQTYDETTLAPDNQYGSEEPQHPGGGQHDQPGTEETESDDIGPATTEVPAIVSTTPEDVVESTIESGYGGAIMTTPPYVDTTTTPKEMQTEATASTKQQPTDSDGNLVDDNGKLLTTTEAPVTTTPVTTAQPTTPTTESPQSSTVTGDGGSTHGQRESTTGGTTIPGTKTPNEEASYATRPTPPADDVCFKAYDDCKKGQPSGGYS*

**>EB_PH2 (accession: PP551272.1)**

MLLSIIFLSLGIVYGQLPDGGDGWELQPDGSYIKTESGGGDSGTGGGGNSGGFYGGGSGGLLGGGSGGLFGGGSGGFLGGGSGGFMSGGNGDWFGGGTWGGGSGGDYGGNGGPPGNGWILQPDGSYMKMETSGGENGGPPGNGWILQPDGTYMKYETGGGGSSHGGGNLPPGFVPPPGFTGNEGGWVLQPDGSYIKTIIETEAPSETIETVNPDYQPDETYSFGKKGSPGYYNIVGFGKPGEMDYFKITPDDQDNPYDYSIEPVVSKDNPSKVTDKNHGRVILRGRKGEPNYEPILQTGKVNRHRYYKIIPDSQKASNYRFMPQRVTGWHTRQGPGEDSGEVLEPEEPEHSEEEPEPIPDEPEIESKNKYVFISLGKPKRYLKLTRHSDNPQDFSLEPVESPDKPDSKPNPNDEEALEVIPQGEKGHPNFKLIVGLGPSYNRRYIQVQPDPNDPNKYHFIPLKSEEQPGEVQTEPNEDDVDFDDIPEDPQYKVIKLPGKTKEHEIFSFGPKGRRIYFKILRRNPNNPNDVEVYPIQDDNEHCESPPSADNPDAPEVVQQGDPNNPDDGPVIKYGPKGKKLLYRLLFKKTGIPKFIPVRQLKKLPGQRRLVFAKQPRIWRFKPQKVKPPIKKPAPQTFETAAPVEETSEYSPDEVDVATEKPESKNPHMILSFGPPKYKVYIKKTPRSPNNPYDYKLEPIGNPQDPNSKPDPKDPNAPEVIETGEIGKPNYAPVVAHGPKSYRRYVIIRPNPKNPNDPNQVEFSPAELVPGSDPKKPKFRVIKAGIPIEPKDPYSESLPKNPKIKIVQKPGDPKKYEVFSFTIKKRRIYIKKVPGATNNPNDFTLEALGNPDDLNSKPDPNDPEAPEIIKGQPIISFGPKGFKKYLIFKPNPQNPKEIQLNPAQLQPGSDPMKPTFIDLVKQILPKPKPVSSAPPVKKEPETKTVPGPAYKIFSVNVKKRRIHLKKIQGPSNNPNDYTLSPLDNPYDPDSKPDPKDPEAPEIIETGQRGTPSFLPILATGPKNKRDYMSVRPSGSGLELRPAEVVPGSDPKHPQFRIPVPGLPKPIIPIPPAAVLPLVLLPGILQSILPVNLGQIQPVIKTYEKPNKKPYTVISFDKKTHIKKTPGPSNNPNDYKLEPIGNPEDPDSKPDPKDPDSPEIIENEKPGTPDYQPVVSFGPVKQRRLITFKVNPSNPKEVHLLPTEIVPGSDPKEHKFRIIPQQQLIPPTKLTDSSELPEVKDVETPDKKHFNIITLRIGPKKIYIKRTPGPSGNPNEYKLEPIGNPDDPNSKPDPKDPDAPEIIESGKPGTPDYQTYVAYGPKPTRKYVSIQKHPSDPNKVAAIPAELEPGSDPKKPRFRIRTLVRPELPVNPLDLLNPLISPFKPHPDIPVPEVKTIIHPDTKKPCHVISIQPKDAKPVFILKTPGPSGSPNDFKLSPLDNPNDVKSTSDPKNPNSPKIFENGKPGTPDYAPIVEYGPKDGKKYVKIAPDNLQPNDPTKLRLTRVKLLLPAKPGEPIKFSELPNVGPKFNPVCYKQYQDCNERS*

**Annotations**

P: hydroxyproline	T/Y/S: phosphorylated residue

C: cysteine	Predicted signal peptide

Underlined: glycine/serine (GS)-rich domain

*Stop codon

Table S6. Predicted Amino acid composition of major high-MW proteins in slime.

| **Residue** | **ER_PH1** | **ER_PH2** | **ES_PH1** | **ES_PH2** | **EB_PH1** | **EB_PH2** |
| --- | --- | --- | --- | --- | --- | --- |
| Ala | 1.5% | 2.4% | 1.9% | 2.6% | 1.7% | 2.3% |
| Arg | 5.4% | 4.9% | 4.8% | 3.6% | 5.6% | 3.2% |
| Asn | 5.0% | 5.6% | 4.5% | 5.7% | 4.3% | 5.7% |
| Asp | 6.7% | 7.8% | 6.8% | 6.6% | 8.0% | 6.4% |
| Cys | 0.2% | 0.3% | 0.3% | 0.2% | 0.4% | 0.3% |
| Gln | 2.0% | 2.9% | 2.9% | 2.9% | 3.5% | 3.4% |
| Glu | 8.2% | 5.4% | 7.2% | 5.8% | 6.3% | 6.3% |
| Gly | 12.7% | 7.5% | 9.0% | 9.6% | 9.6% | 9.7% |
| His | 3.0% | 2.4% | 2.9% | 1.6% | 3.0% | 1.5% |
| Ile | 3.3% | 5.9% | 3.7% | 6.0% | 3.5% | 6.7% |
| Leu | 2.4% | 4.2% | 3.2% | 5.7% | 3.1% | 5.6% |
| Lys | 10.1% | 11.3% | 10.2% | 9.2% | 9.7% | 9.9% |
| Met | 0.3% | 0.6% | 0.6% | 0.7% | 0.5% | 0.6% |
| Phe | 2.9% | 3.3% | 3.4% | 3.0% | 2.8% | 3.2% |
| Pro | 18.0% | 16.7% | 17.7% | 16.0% | 16.7% | 15.6% |
| Ser | 3.6% | 5.3% | 5.2% | 6.7% | 3.9% | 5.8% |
| Thr | 6.0% | 4.0% | 6.7% | 4.1% | 7.4% | 4.0% |
| Trp | 0.1% | 0.4% | 0.1% | 0.4% | 0.2% | 0.5% |
| Tyr | 3.7% | 3.4% | 3.4% | 3.9% | 4.0% | 3.9% |
| Val | 5.0% | 5.6% | 5.6% | 5.6% | 5.8% | 5.3% |

Table S7. Single nucleotide and indel polymorphism. Identified in high-MW proteins in re-sequencing comparing to the sequences obtained from RNA-sequencing and resulted amino acid composition changes.

| **Protein** | **Position** | **RNAseq** | **Amino acid** | **Re-seq** | **Amino acid** |
| --- | --- | --- | --- | --- | --- |
| ER_PH1 | 644 | AAA | K | AGA | R |
|  | 672 | GAA | E | AAA | K |
|  | 695 | AGG | R | AAT | N |
|  | 712 | AAG | K | AGT | S |
|  | 724 | GAA | E | GAT | D |
|  | 736 | ACA | T | CCA | P |
|  | 742 | GCC | A | ACC | T |
|  | 774 | ATT | I | ACT | T |
|  | 787 | CCT | P | GCT | A |
|  | 794 | GCT | A | TTT | F |
|  | 807 | GAC | D | GGC | G |
|  | 817 | GTA | V | ATA | I |
|  | 847 | AAA | K | AGA | R |
|  | 848 | CAT | H | CAG | Q |
|  | 850 | AGG | R | AGT | S |
|  | 866 | GCT | A | ATA | I |
|  | 878 | TTT | F | TGT | C |
|  | 891 | AAT | N | CAT | H |
|  | 895 | GAA | E | CAA | Q |
|  | 909 | TAC | Y | AAC | N |
|  | 927 | CTT | L | ATT | I |
|  | 934 | AAA | K | CAT | H |
|  | 938 | AAA | K | GAA | E |
|  | 953 | CCC | P | TCC | S |
|  | 955 | GGT | G | / | / |
|  | 957 | GAT | D | AAT | N |
|  | 961 | CGT | R | TGT | C |
|  | 964 | ACT | T | AAT | N |
|  | 1000 | GAT | D | AGT/GGT | S/G |
|  | 1001 | GGT | G | AGT | S |
|  | 1007 | GAC | D | GAA | E |
|  | 1008 | AAA | K | ATA | I |
|  | 1026 | GCT | A | ACT/TCT | T/S |
|  | 1038 | GTA | V | ATA | I |
|  | 1054 | CAT | H | CGT | R |
|  | 1069 | CAA | Q | ACA | T |
|  | 1079 | CGT | R | CAT | H |
|  | 1091 | AAT | N | AAG | K |
|  | 1093 | CCA | P | TCA | S |
|  | 1121 | GAT | D | AAT | N |
|  | 1127 | GAA | E | CAA | Q |
|  | 1173 | CGG | R | CAA | Q |
|  | 1184 | GAA | E | GTA | V |
|  | 1199 | AGA | R | AAA | K |
|  | 1203 | CAA | Q | CGA | R |
|  | 1235 | AAA | K | ACA | T |
|  | 1247 | CCT | P | CAA | Q |
|  | 1262 | ATA | I | ACA | T |
|  | 1284 | AAA | K | ACA | T |
|  | 1293 | AAT | N | AAA | K |
|  | 1302 | GAA | E | GTA | V |
|  | 1330 | / | / | ACA | T |
|  | 1356 | GAC | D | GAG | E |
|  | 1401 | AGA | R | AAA | K |
| ER_PH2 | 441 | / | / | GCA | A |
|  | 479 | ATA | I | ACA | T |
|  | 484 | AAA | K | CAA | Q |
|  | 495 | AGT | S | CGT | R |
|  | 509 | CTA | L | CCA | P |
|  | 510 | TCA | S | GCA | A |
|  | 551 | GTC | V | ATC | I |
|  | 574 | ACT | T | AAT | N |
|  | 579 | GAG | E | CAA | Q |
|  | 597 | GAT | D | GAA | E |
|  | 603 | GAA | E | GAT | D |
|  | 608 | CAT | H | GGT | G |
|  | 611 | GAT | D | GGT | G |
|  | 631 | CCA | P | CAA | Q |
|  | 639 | GTT | V | TTT | F |
|  | 640 | AAA | K | TTA | L |
|  | 669 | AGA | R | AAA | K |
|  | 671 | AAG | K | GAG | E |
|  | 672 | CCC | P | CTC | L |
|  | 682 | AAT | N | GAT | D |
|  | 683 | TTT | F | TAT | Y |
|  | 684 | ACA | T | AAA | K |
|  | 686 | CAT | H | CTT | L |
|  | 697 | ATT | I | GTT | V |
|  | 701 | ATA | I | AGA | R |
|  | 706 | CCA | P | ACA | T |
|  | 707 | TTC | F | CAC | H |
|  | 708 | TCC | S | GCC | A |
|  | 709 | CAT | H | ATG | M |
|  | 710 | GTT | V | CTT | L |
|  | 711 | CGC | R | GAC | D |
|  | 712 | TTC | F | TAC | Y |
|  | 715 | GTA | V | GCT | A |
|  | 718 | AAA | K | ACA | T |
|  | 719 | CGC | R | CCC | P |
|  | 720 | AAA | K | AAT | N |
|  | 722 | GAG | E | AAG | K |
|  | 723 | CAA | Q | CCA | P |
|  | 725 | TCT | S | CAT | H |
|  | 728 | ATA | I | TTA | L |
|  | 789 | CCA | P | CTA | L |
|  | 793 | GAT | D | AAT | N |
|  | 799 | ATA | I | GTA | V |
| EB_PH1 | 304 | CAA | Q | CCA | P |
|  | 517 | GAC | D | AAT | N |
|  | 519 | AAA | K | CGA | R |
|  | 520 | GAT | D | AAT | N |
|  | 524 | AGA | R | AGT | S |
|  | 569 | ACT | T | AAT | N |
|  | 570 | CAT | H | CGT | R |
|  | 612 | ATT | I | ACT | T |
|  | 619 | GAA | E | GAT | D |
|  | 627 | ACC | T | CCA | P |
|  | 630 | AAA | K | AAC | N |
|  | 631 | CCA | P | CGA | R |
|  | 637 | AGA | R | AAA | K |
|  | 644 | ATA | I | CTA | L |
|  | 645 | GGA | G | GAA | E |
|  | 646 | AAT | N | GAT | D |
|  | 647 | AAA | K | GAG | E |
|  | 649 | CAT | H | TAT | Y |
|  | 650 | CAA | Q | CGA | R |
|  | 653 | TTA | L | TTT | F |
|  | 1516 | TCT | S | TAT | Y |
| EB_PH2 | 1063 | ACA | T | AAA | K |
|  | 1330 | CAA | Q | AAA | K |
|  | 1346 | GAT | D | GAA | E |
|  | 1350 | GTT | V | GAT | D |
|  | 1372 | GTT | V | GAT | D |
|  | 1421 | CAT | H | AAA | K |

Table S8. Re-sequencing primers for high-MW constructs.

| **Construct** | **Gene-specific primers (5’**–**3’)** |
| --- | --- |
| ER_PH1 | **GSP-1**: GATTACGCCAAGCTTTGGGCGCCCTGATTTTCCTGGTCT  **GSP-2**: GATTACGCCAAGCTTCCGTCCTGCTTCCACTCCAGAGCC |
| ER_PH2 | **GSP-1**: GATTACGCCAAGCTTACCGATTGGCGGTTTATGGGGCT  **GSP-2**: GATTACGCCAAGCTTAGAGCCCCTTGGCAACCCTGA |
| EB_PH1 | **GSP-1**: GATTACGCCAAGCTTAGCGTTGTGGATGCTCCTTTGG  **GSP-2**: GATTACGCCAAGCTTACCAGGCCGTCAGGGAAAGCCAG  ***GSP-Int1**: GATTACGCCAAGCTTCCAAAGGAGCATCCACAACGCT  ***GSP-Int2**: GATTACGCCAAGCTTCTGGCTTTCCCTGACGGCCTGGT |
| EB_PH2 | **GSP-1**: GATTACGCCAAGCTTTGCCACCCAGTCACACGCTGAGG  **GSP-2**: GATTACGCCAAGCTTAGACCAGCAGAGGTAGTCCCTGGT  ***GSP-Int1**: GATTACGCCAAGCTTCCTCAGCGTGTGACTGGGTGGCA  ***GSP-Int2**: GATTACGCCAAGCTTACCAGGGACTACCTCTGCTGGTCT |

*The primers were used for amplification of regions not covered by 5’/3’ RACE PCR.

**Table S9.** Molecular sequences used for phylogenetic analyses with corresponding GenBank accession numbers.

| **Species** | **Accession number (by gene)** | | |
| --- | --- | --- | --- |
|  | ***COI*** | ***12S rRNA*** | ***16s rRNA*** |
| **Peripatidae** | | | |
| *Eoperipatus* sp. (Singapore) | present study –  to be added | present study –  to be added | present study –  to be added |
| *Eoperipatus* sp. (Thailand) | KY322403 | JX568982 | KY322424 |
| *Epiperipatus* cf. *barbadensis* (Froehlich, 1962) | present study –  to be added | present study –  to be added | present study –  to be added |
| *Epiperipatus edwardsii* (Blanchard, 1847) | HG531958 | HG531961 | HG531962 |
| *Epiperipatus vagans* (Brues, 1925) | MH107347 | MG973666 | MG973485 |
| *Mesoperipatus tholloni* (Bouvier, 1898) | KC754645 | KC754478 | KC754528 |
| *Oroperipatus eisenii* (Wheeler, 1898) | MH107369 | MG973712 | MG973531 |
| *Plicatoperipatus jamaicensis* (Grabham & Cockerell, 1892) | KC754639 | MG973674 | MG973506 |
| *Principapillatus hitoyensis* Oliveira *et al.*, 2012 | KC754642 | KC754476 | KC754525 |
| **Peripatopsidae** | | | |
| *Diemenipatus mesibovi* Oliveira *et al.*, 2018 | MG692748 | MG692758 | MG692768 |
| *Diemenipatus taiti* Oliveira *et al.*, 2018 [locality I] | MG692746 | MG692756 | MG692766 |
| *Diemenipatus taiti* Oliveira *et al.*, 2018 [locality II] | MG692747 | MG692757 | MG692767 |
| *Euperipatoides leuckartii* (Sänger, 1871) | KC754649 | KC754481 | KC754531 |
| *Euperipatoides rowelli* Reid, 1996 | KY322408 | KY322435 | KY322429 |
| *Kumbadjena occidentalis* (Fletcher, 1895) | KC754653 | KC754484 | KC754535 |
| *Leucopatus anophthalmus* (Ruhberg *et al.*, 1991) | KC754691 | KC754517 | KC754572 |
| *Metaperipatus blainvillei* (Gervais, 1837) | KR907171 | KR906981 | KC754540 |
| *Metaperipatus inae* Mayer, 2007 | KC754659 | KC754490 | KC754541 |
| *Occiperipatoides gilesii* (Spencer, 1909) | KC754660 | KC754491 | KC754542 |
| *Ooperipatellus cryptus* Jackson & Taylor, 1994 | KC754662 | KC754492 | KC754544 |
| *Ooperipatellus decoratus* (Baehr, 1977) | KY322407 | KY322434 | KY322428 |
| *Ooperipatellus insignis* (Dendy, 1890) | KY322406 | KY322433 | KY322427 |
| *Ooperipatellus nanus* Ruhberg, 1985 | KC754665 | KC754495 | KC754547 |
| *Ooperipatellus nickmayeri* Oliveira & Mayer, 2017 | KY322409 | KY322436 | KY322430 |
| *Ooperipatus birrgus* Reid, 2000 | KC754671 | KC754501 | KC754553 |
| *Ooperipatus caesius* Reid, 2000 | KC754672 | KC754502 | KC754554 |
| *Ooperipatus porcatus* Reid, 2000 | KC754673 | KC754503 | KC754555 |
| *Peripatoides* sp. 3 | KC754681 | KC754509 | KC754562 |
| *Peripatoides* sp. 4 | KC754682 | KC754510 | KC754563 |
| *Peripatoides novaezealandiae* (Hutton, 1876) I | KC754677 | KC754506 | KC754559 |
| *Peripatoides novaezealandiae* (Hutton, 1876) II | KC754684 | KC754512 | KC754565 |
| *Peripatoides suteri* (Dendy, 1894) | KC754683 | KC754511 | KC754564 |
| *Peripatoides sympatrica* Trewick, 1998 | KC754684 | KC754512 | KC754565 |
| *Peripatopsis capensis* (Grube, 1866) | KC754685 | KC754513 | KC754566 |
| *Peripatopsis lawrencei* McDonald *et al.*, 2012 | KC754687 | KC754514 | KC754568 |
| *Peripatopsis moseleyi* (Wood-Mason, 1879) | KC754688 | KC754515 | KC754569 |
| *Tasmanipatus barretti* Ruhberg *et al.*, 1991 | KC754692 | KC754518 | KC754573 |

Movie S1 (separate file). Rotating structure of the mid-MW LRR protein ER_PM1-var1 in slime of *Eu. rowelli*. 1) Model confidence, highest confidence in dark blue, high confidence in light blue, low confidence in yellow, lowest confidence in orange; 2) Annotated secondary conformations, α-helices in red, β-sheets in green; 3) Surface charges, positively charged residues in red, negatively

Movie S2 (separate file). Rotating structure of the high-MW LRR protein ER_PH1 in slime of *Eu. rowelli*. 1) Model confidence, highest confidence in dark blue, high confidence in light blue, low confidence in yellow, lowest confidence in orange; 2) Annotated secondary conformations, α-helices in red, β-sheets in green; 3) Surface charges, positively charged residues in red, negatively charged residues in blue; 4) Surface hydrophobicity, gradient from hydrophilic in white to hydrophobic in red.
